# Stalling of the endometrial decidual reaction determines the recurrence risk of miscarriage

**DOI:** 10.1101/2024.11.07.622412

**Authors:** Joanne Muter, Chow-Seng Kong, Mireia Taus Nebot, Maria Tryfonos, Pavle Vrljicak, Paul J. Brighton, Danai B. Dimakou, Megan Vickers, Hiroyuki Yoshihara, Sascha Ott, Bee K. Tan, Phillip R. Bennett, Siobhan Quenby, Alex Richter, Hilde Van de Velde, Emma S. Lucas, Thomas M. Rawlings, Jan J. Brosens

## Abstract

In each menstrual cycle, progesterone acting on estrogen-primed endometrium elicits an inflammatory decidual reaction, rendering it poised for embryo implantation and transformation into the decidua of pregnancy. Here, we show that the sequential functions of the decidual reaction - implantation and decidualization - pivot on the time-sensitive loss of progesterone-resistant stromal cells that form a transient implantation niche and reciprocal expansion of progesterone-responsive pre-decidual cells. In parallel, proliferation and differentiation increase the abundance of immunotolerant uterine natural killer (uNK) cells. Examination of pre-pregnancy endometrial biopsies from 924 women revealed that the frequency of cycles culminating in a blunted or stalled decidual reaction closely aligns with the age-independent recurrence risk of miscarriage. Further, analysis of 632 biopsies obtained in different cycles from 316 women indicated that prior miscarriages disrupt intercycle endometrial homeostasis, an observation supported by modelling the impacts of prolonged decidual inflammation in three-dimensional endometrial assembloids. Although stalling of the decidual reaction is often accompanied by a poor expansion of immunotolerant uNK cells, miscarriages do not impact intercycle uNK cell dynamics. Our findings indicate that intrinsic uterine mechanisms hardwire the recurrence risk of miscarriage, underscoring the need for pre-pregnancy diagnostics and therapeutics.

**One Sentence Summary:** The frequency of menstrual cycles culminating in a suboptimal decidual reaction determines the recurrence risk of miscarriage.

## INTRODUCTION

Miscarriage denotes the loss of a pregnancy before viability (*1*). Approximately one in three embryos perish following implantation in healthy women, although often before routine detection of pregnancy (*2, 3*). This attrition rate reflects the high prevalence of chromosomal errors in preimplantation human embryos (*4*) and the physiological role of the endometrium in selecting against low-fitness embryos (*2, 3, 5, 6*). The pooled miscarriage risk in all clinically recognized pregnancies is an estimated 15% (*1*), with most losses (∼90%) occurring before the onset of uteroplacental perfusion at the end of the first trimester (*7*). Epidemiological studies consistently highlight that two factors, maternal age and the number of preceding pregnancy losses, disproportionally impact miscarriage rates (*7–9*). The age-dependent risk reflects the increase in aneuploid pregnancies in women aged 35 years and older, mirroring the incidence of meiotic chromosome errors in oocytes and embryos (*10, 11*). Each prior pregnancy loss further compounds the risk stepwise by 5-10% (*7–9*), but the underlying mechanism is unknown. A plausible but untested hypothesis is that the recurrence risk of miscarriage reflects the frequency of menstrual cycles culminating in an endometrial environment permissive of embryo implantation but inadequately prepared for decidual transformation (*3*), that is, the formation of a robust immunotolerant matrix that anchors and supports the semi-allogenic placenta throughout pregnancy (*2, 5*).

Each menstrual cycle starts with the shedding of the superficial endometrial layer, bleeding, and re-epithelization of the basal layer (*2*). Following menstruation, estradiol-dependent regeneration of the superficial layer, on average, quadruples the thickness and volume of the uterine mucosa prior to ovulation (*12*). Local morphogen and cytokine gradients regulate epithelial and stromal cell proliferation, resulting in tissue stratification and positional cell specification (*2*). After ovulation, progesterone acting on this spatial template triggers a decidual reaction, an endogenous inflammatory tissue response that heralds the start of the four-day midluteal implantation window (*2, 13*). Histologically, the implantation window coincides with the onset of glandular secretion, marked oedema, and proliferative expansion and differentiation of uterine natural killer (uNK) cells (*14, 15*). A canonical feature of the decidual reaction is the progesterone-dependent reprogramming of stromal cells, termed pre-decidual cells, under the control of evolutionarily conserved transcription factors (*5, 16, 17*). By secreting IL-15, pre-decidual cells activate cytolytic uNK cells to prune stressed and damaged cells, curtailing tissue inflammation (*18, 19*). The emergence of mature decidual cells, characterized by their epithelioid morphology and stress-resistant phenotype, signals the window’s closure (*5, 14*).

Implantation in humans is interstitial, meaning the embryo fully breaches the luminal epithelium and embeds in the underlying stroma (*2, 20*). Modelling this process in endometrial assembloids, comprising gland-like organoids and primary stromal cells, showed that the decidual reaction imposes a broad binary state on stromal cells, separating progesterone-responsive and -resistant subsets (*21*). Progesterone-resistant stromal cells, including transitional epithelial-mesenchymal and acutely senescent cells, create a dynamic implantation environment for co-cultured embryos. However, their persistence also leads to spontaneous disintegration of assembloids. By contrast, the lack of progesterone-resistant stromal cells accelerates the emergence of decidual cells and entraps embryos in a static matrix, precluding implantation (*21*). These observations suggest that implantation and decidual transformation require time-sensitive rebalancing of functionally distinct stromal subsets across the implantation window. Here, we characterize this process *in vivo*, explore the interplay between stromal and uNK cell dynamics, and examine if the frequency of menstrual cycles resulting in a blunted or stalled decidual reaction accounts for the recurrence risk of miscarriage.

## RESULTS

### Characterization of *DIO2*+ and *PLA2G2A*+ endometrial stromal subsets

We previously reported that *SCARA5* (ferritin receptor) and *DIO2* (iodothyronine deiodinase 2), genes induced and repressed by progesterone, respectively, mark different stromal subsets (*19*). When normalized for the timing of an endometrial sample relative to the pre-ovulatory luteinizing hormone (LH) surge, high endometrial *DIO2* expression mirrors lower *SCARA5* transcript levels and vice versa (*19*), indicating that the ratio of these marker genes is a putative measure of the overall progesterone responsiveness of stromal cells in tissue samples. However, *SCARA5* is expressed across the luteal phase in a relatively narrow dynamic range (*19*), suggesting it lacks sensitivity to monitor emerging pre-decidual cells. To explore this further, we measured *SCARA5* and *DIO2* transcripts in twelve endometrial biopsies obtained in different menstrual cycles from six subjects (Table S1). Following normalization to the LH surge (*19*), progesterone responsiveness was deemed normal in nine samples (*SCARA5*/*DIO2* ≥ 25^th^ percentile) and impaired in three biopsies (*SCARA5*/*DIO2* < 25^th^ percentile) (Fig. 1A). RNA-sequencing revealed that high *DIO2* transcript levels in the three biopsies designated as progesterone-resistant were associated with loss of *PLA2G2A*, much more so than *SCARA5* (Fig. 1B and Data S1). *PLA2G2A* encodes the acute-phase protein phospholipase A2 group IIA (PLA2G2A). By hydrolyzing the ester bond of the fatty acyl group attached at the sn-2 position of phospholipids, PLA2G2A promotes the release of arachidonic acid (*22*), the first step in the production of the ancestral decidual signal, prostaglandin E2 (*23*). Multiplexed single molecule *in situ* hybridization (smISH) demonstrated that *PLA2G2A* expression marks stromal cells separated from the luminal epithelium by a layer of *DIO2*+ cells (Fig. 1C). To characterize the *DIO2*+ and *PLA2G2A*+ subsets, we performed single-cell RNA-sequencing (scRNA-seq) on twelve additional biopsies timed to span the implantation window (Table S2). Following dimensionality reduction, seven major cell clusters were identified, including 33,416 stromal cells (Fig. 1D, fig. S1A and Data file S1). Expression of *PLA2G2A*, like *DIO2*, was highly enriched in stromal cells (*p* < 2.3 × 10^-308^, Fig. 1B).

**Fig. 1.**
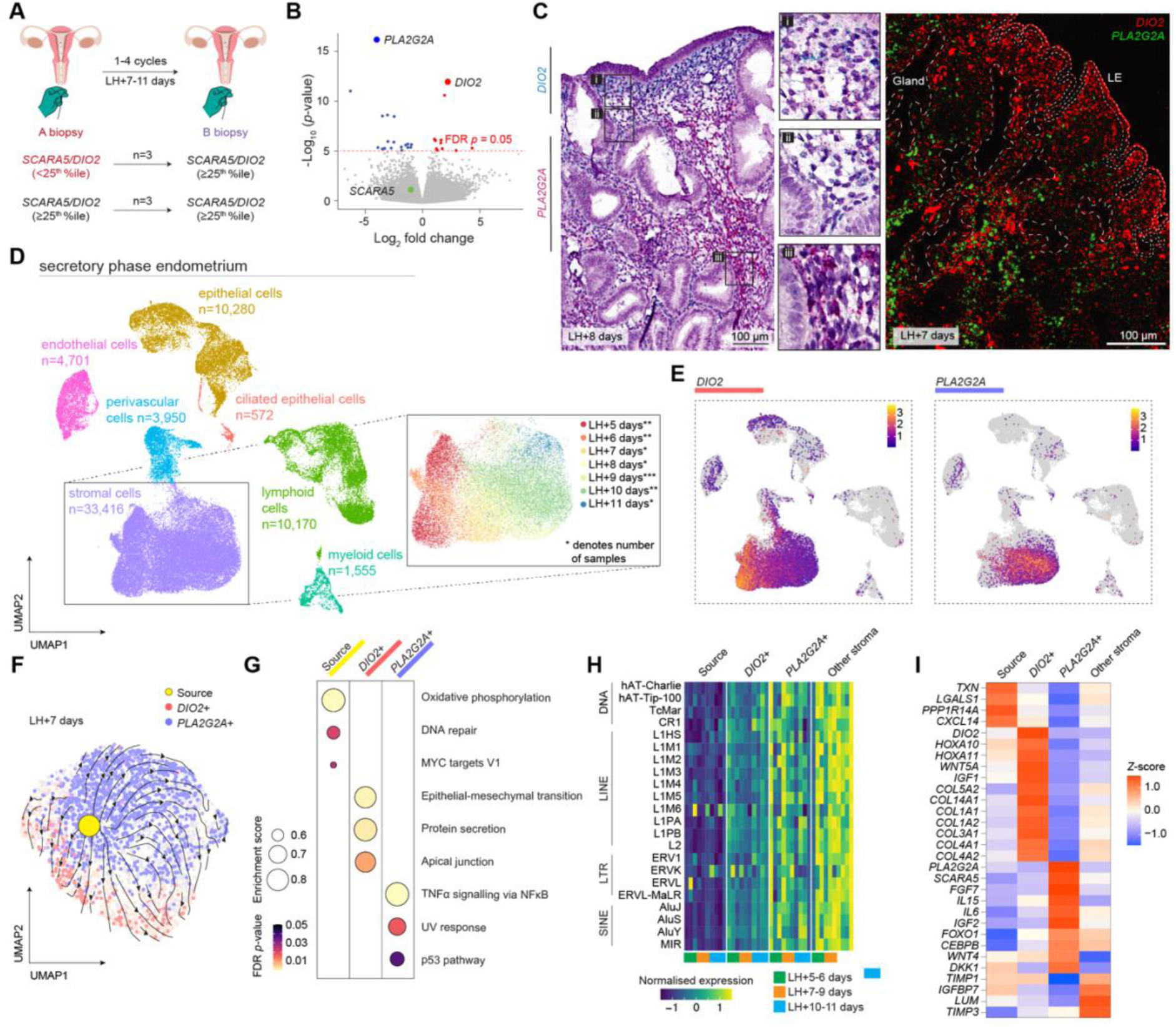
*DIO2*+ and *PLA2G2A*+ mark functionally distinct stromal subsets. (**A**) Depiction of endometrial sampling in six subjects across two menstrual cycles. For each sample (n=12), the ratio of *SCARA5* and *DIO2* transcripts measured by RT-qPCR was normalized for the day of the biopsy relative to the LH surge; samples were deemed progesterone-resistant (*SCARA5/DIO2* < 25%ile, highlighted in red) or progesterone-responsive (*SCARA5/DIO2* ≥ 25%ile). (**B**) Volcano plot showing gene expression variance between progesterone-resistant and -responsive endometrial biopsies. Transcripts of interest are highlighted. (**C**) Representative images of *DIO2* and *PLA2G2A* transcripts in midluteal endometrium by chromogenic (left) and fluorescent (right) smISH. LE, luminal epithelium. (**D**) UMAP projections of scRNA-seq data showing the cellular composition of luteal phase endometrium (n=12). The color key in the inset depicts days post-LH surge; asterisks denote sample numbers for each day. (**E**) Feature plots showing relative expression of *DIO2* and *PLA2G2A* in different cell types. The colour scale represents log-transformed average gene expression. (**F**) RNA velocity stream mapped onto a UMAP plot of stromal subsets from a midluteal biopsy. (**G**) GSEA enrichment plot. The dot size corresponds to the enrichment score; the color key denotes Benjamini-Hochberg-adjusted *p*-values. (**H**) Heatmap of normalized transposable element (TE)-derived transcript levels encompassing DNA transposons (DNA), long interspersed nuclear elements (LINE), long terminal repeats retrotransposons (LTR), and short interspersed nuclear elements (SINEs). (**I**) Heatmap of z-score-scaled genes in source, *DIO2*+, *PLA2G2A*+ and other stromal cells.

Temporal changes in gene expression were evident upon clustering of stromal cells by the day of the biopsy relative to LH surge (LH+5 to +11 days) (Fig. 1D). Further, the putative implantation window (LH+6 to +9 days) coincided with a conspicuous shift from a preponderance of *DIO2*+ to an abundance of *PLA2G2A*+ cells (Fig. 1E and fig. S1C). *DIO2*+ and *PLA2G2A*+ subsets constituted ∼43% of all captured stromal cells, but cells co-expressing both genes were relatively rare (Fig. 1C), averaging at 3.6% (SD: ± 2.9%) (fig. S1C). In ten out of twelve samples, RNA velocity analysis inferred that a discrete cluster of source cells drives gene expression in the *DIO2*+ and *PLA2G2A*+ subsets (Fig. 1F and fig. S1D). Hallmark gene set enrichment analysis (GSEA) predicted heightened oxidative phosphorylation and activation of DNA repair pathways in source cells (Fig. 1G), indicative of acute cellular stress. Notably, source cells, and to a lesser extent *DIO2*+ cells, differ from other stromal subsets by the depletion of transcripts derived from transposable elements (TEs) (Fig. 1H), suggesting lack of prior exposure to replication stress (*24*). Visualization of hallmark DNA repair genes on a spatial transcriptomic map of midluteal endometrium identified putative ‘hotspots’ of damaged cells residing some distance from the luminal epithelium (fig. S1E). Decidual cells may have emerged in ancestral eutherians due to progesterone gaining control over the senescence-associated secretory phenotype (SASP), converting a pro-inflammatory cell state into an anti-inflammatory state (*17*). In keeping with this paradigm, the enriched hallmark gene sets in *PLA2G2A*+ cells predicted NF-κB and p53 activation (Fig. 1G), two pivotal signaling pathways in cellular senescence (*25*). Transcriptional profiling confirmed that *PLA2G2A*+ cells are pre-decidual cells, exemplified by the expression of genes coding canonical decidual transcription factors (e.g., *FOXO1* and *CEBPB*) and secreted proteins (e.g., *IL15*, *WNT4*, *IGF2*, *FGF7* and *DKK1*) (Fig. 1I) (*5*). By contrast, *DIO2*+ cells are enriched in hallmark epithelial-mesenchymal transition (EMT) genes (Fig. 1G). DIO2 regulates energy expenditure in cells by catalyzing the conversion of prohormone thyroxine (3,5,3’,5’-tetraiodothyronine, T4) to the bioactive thyroid hormone (3,5,3’-triiodothyronine, T3)(*26*). Subluminal *DIO2*+ cells produce an abundance of extracellular matrix (ECM) components (Fig. 1I), plausibly explaining the need for heightened energy expenditure. *DIO2*+ cells further express multiple genes that govern Müllerian duct patterning during development, including *WNT5A*, *HOXA10* and *HOXA11* (Fig. 1I) (*27*). Thus, an embryo breaching the luminal epithelium encounters a specialized stromal microenvironment harboring *DIO2*+ cells. The EMT gene signature predicts that subluminal *DIO2*+ cells have a migratory and invasive phenotype, properties considered essential for interstitial implantation (*5, 6*). Underlying this specialized implantation niche are the precursors of anti-inflammatory decidual cells, which in pregnancy cooperate with invasive trophoblast and local immune cells to form the uteroplacental interface (*28, 29*).

### Spatiotemporal regulation of uNK subsets

Accumulation of uNK cells in the endometrium is a consistent feature of the decidual reaction in menstruating primates (*30*). NK cells are effector lymphocytes of the innate immune system involved in the clearance of stressed and damaged cells, allorecognition and antimicrobial defenses (*31*). In human endometrium, proliferative expansion promotes differentiation of CD56+ uNK cells, characterized by the sequential acquisition of killer cell immunoglobulin-like receptors (KIRs) and the ectonucleotidase CD39 (*15*). As local cues regulate tissue-resident NK cells (*15*), we examined the spatial distribution of CD56+KIR-and CD56+KIR+ uNK cells during the implantation window. Dual-color fluorescence microscopy revealed an overrepresentation of CD56+KIR+ uNK cells in stroma abutting the luminal epithelium and CD56+KIR-uNK cells in the underlying tissue (Fig. 2A). Flow cytometric analysis of 55 timed biopsies (Table S2) confirmed the rapid expansion of KIR+CD39-and KIR+CD39+ uNK subsets as the endometrium cycles across the implantation window (Fig. 2, B and C, fig. S2A).

**Fig. 2.**
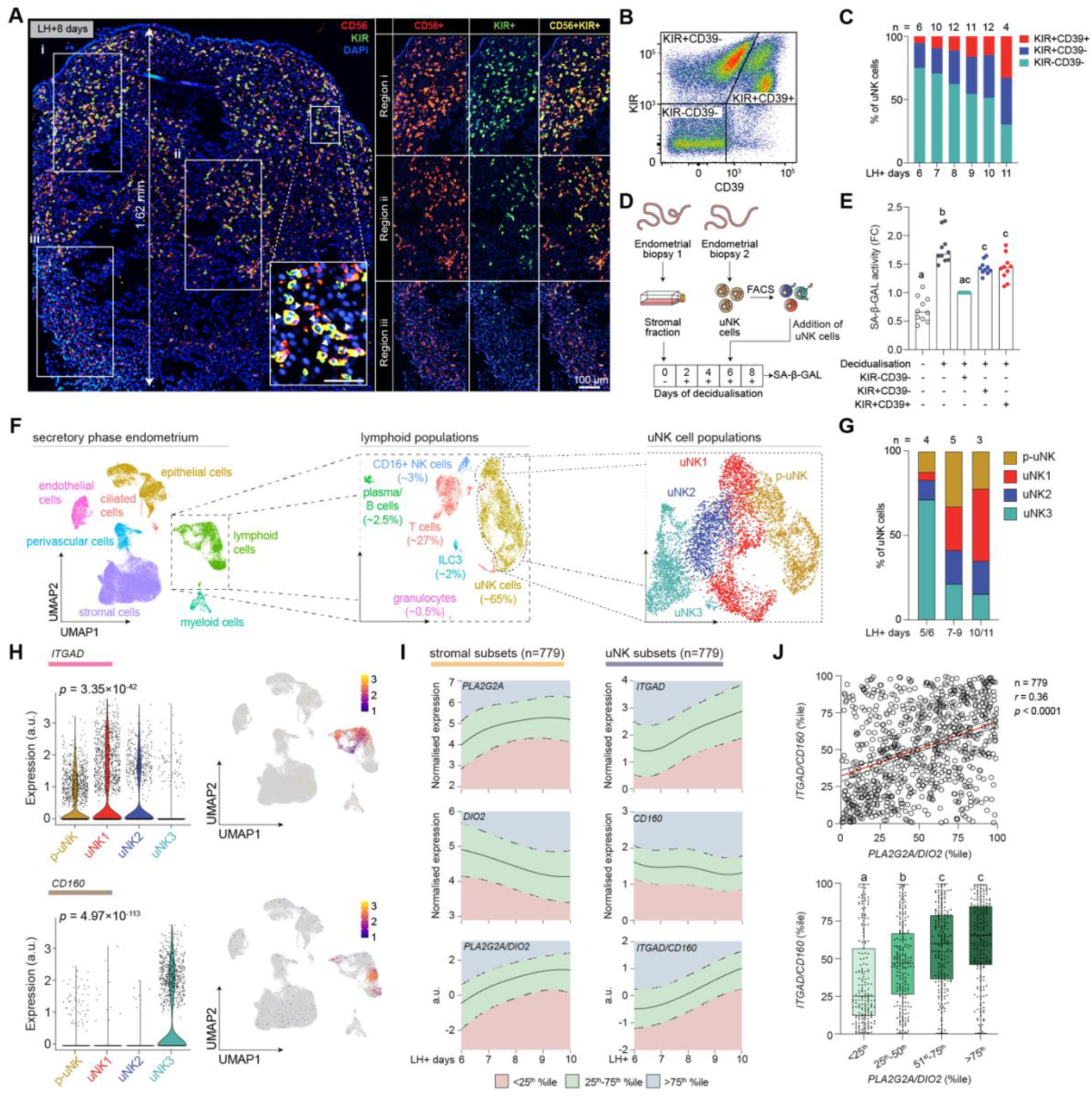
Spatiotemporal dynamics of uNK cells. (**A**) Representative image of CD56 (NCAM1) and pan-KIR immunofluorescence in midluteal endometrium. CD56 and pan-KIR immunoreactivity in different regions is shown separately in boxes A-C. The insert shows a region at higher magnification. Scale bar = 50 µm. Nuclei are stained with DAPI. (**B**) Flow cytometry density plot showing different uNK subsets based on KIR and CD39 expression. (**C**) Bar graph depicting the temporal changes in the relative abundance of uNK subsets in 55 endometrial samples as determined by flow cytometry. (**D**) Schematic of experimental design. Primary stromal cultures first decidualized for six days were co-cultured with different uNK subsets for two additional days. The uNK subsets were purified by FACS using pan-KIR and CD39 antibodies. (**E**) Bar graphs showing median fold change (FC) in SA-β-GAL activity in 10 biological repeat experiments. Individual data points are also shown. Different letters indicate statistically significant differences at *p* < 0.01, Friedman test with Dunn’s multiple comparisons test. (**F**) UMAP projections of scRNA-seq data showing lymphoid populations and uNK subsets. (**G**) Bar graphs depicting the temporal changes in the relative abundance of uNK subsets as determined by scRNA-seq. (**H**) Violin plots (left) and feature plots (right) of *ITGAD* and *CD160* expression with *p*-values based on Wilcoxon rank sum test; a.u., arbitrary units. (**I**) Relative expression of stromal and uNK subset marker genes and their ratios in 779 endometrial biopsies obtained 6 to 10 days after the LH surge. The solid line indicates median expression; the dotted lines mark the boundaries of the upper and lower quartiles. Ratios were fitted against a gamma or a log-normal distribution depending on LH+day; a.u., arbitrary units. (**J**) Upper panel: Spearman’s correlation between *ITGAD/CD160* and *PLA2G2A/DIO2* %iles. Lower panel: box plots comparing *ITGAD/CD160* %iles and *PLA2G2A/DIO2* %iles grouped in quartile bins. Different letters above the whiskers indicate significance between groups at *p* < 0.05, Kruskal-Wallis test with Dunn’s multiple comparisons test.

We reasoned that the spatial organization of phenotypically distinct uNK cells reflects functional adaptation to specific stromal microenvironments. We investigated this possibility by co-culturing decidualizing primary stromal cells with CD56+ uNK cells separated first by fluorescence-activated cell sorting (FACS) into different subsets based on the expression of KIR and CD39 (Fig. 2D and Table S2). Senescence-associated β-galactosidase (SA-β-GAL) activity increases markedly upon decidualization (Fig. 2E), reflecting the emergence of decidual-like senescent cells (*18, 19*). Co-cultured uNK cells attenuated SA-β-GAL activity with KIR-CD39-uNK cells being twice as effective than KIR+CD39-or KIR+CD39+ uNK cells (Fig. 2E). These observations imply that KIR-CD39-uNK cells, which reside with *PLA2G2A*+ pre-decidual cells, are cytolytic cells that constrain decidual inflammation through targeted killing of stressed and damaged cells. The crowding of attenuated KIR+ uNK cells in the subluminal implantation niche is in keeping with their known roles in pregnancy, including conferring maternal immune tolerance to the semi-allogenic conceptus and promoting trophoblast invasion (*28*).

Based on scRNA-seq analysis, uNK cells comprise ∼65% of lymphoid immune cells in the midluteal endometrium, more than two-fold the abundance of T cells (∼27%) (Fig. 2F, fig. 2B and Data file S1). Apart from a discrete cluster of peripheral blood CD16+ NK cells (∼3.6% of lymphoid cells), we identified four uNK subsets (Fig. 2F), in keeping with other studies (*15, 32*). All *KIR*+ uNK subsets (uNK1, uNK2 and proliferating uNK cells) expand rapidly upon opening of the implantation window, thus diluting the relative abundance of uNK3 cells, corresponding to cytolytic KIR-CD39-uNK cells (Fig. 2G and fig. S2C). We identified *ITGAD* (integrin subunit alpha D) and *CD160* (CD160 molecule) as genes selectively enriched, respectively, in *KIR*+ and *KIR*-uNK subsets (Fig. 2H and fig. S2D). In agreement, the ratio of *ITGAD*/*CD160* transcript levels discriminated between freshly isolated KIR-and KIR+ uNK cells (fig. S2E). To gain further insights into the cellular dynamics across the implantation window, we quantified the expression of genes marking the stromal (*PLA2G2A*, *DIO2*) and uNK (*ITGAD*, *CD160*) subsets in 779 timed endometrial samples (Table S2). Next, we calculated the ratios of paired marker genes and established percentile ranks (Fig. 2I). This analysis revealed a significant correlation (*r* = 0.36, *p* < 0.0001) between the strength of the decidual reaction in stromal cells, as measured by *PLA2G2A*/*DIO2* percentiles, and magnitude of uNK cell expansion (*ITGAD*/*CD160* percentiles) (Fig. 2J). However, divergence was apparent in a substantial proportion of samples, suggesting that time-sensitive rebalancing of stromal and uNK subsets are coregulated but not necessarily interdependent processes.

### Regulation of stromal and uNK subsets by hormonal and embryonic cues

Embryos implant preferentially near the uterine fundus (*20*), raising the possibility that spatial differences preclude accurate assessment of the decidual reaction or uNK cell expansion in tissue samples. Pipelle biopsies capture the superficial endometrium at 1-2 mm depth, but samples can be several centimeters long (Fig. 3A). Analysis of marker genes across the entire length of biopsies or at opposite ends of the samples (n=264) showed that spatial variability is limited, albeit more pronounced for uNK than stromal subsets (Fig. 3, B to E and Table S2). Next, we measured circulating progesterone, estradiol, and thyroid-stimulating hormone (TSH) levels in 315 subjects (Table S2). Progesterone and estradiol levels, normalized to the LH surge (fig. S3A), correlated positively with *PLA2G2A*/*DIO2* but not *ITGAD*/*CD160* ratios (Fig. 3F), reflecting the direct hormonal dependency of pre-decidual cells, but not uNK cells (*5, 15*). Although uNK cells originate from circulating progenitor cells (*15*), we observed no correlation between the abundance of peripheral blood NK cells and local uNK cell expansion as measured by normalized *ITGAD*/*CD160* ratios (fig. S3B). TSH levels had no discernible impact on stromal or uNK subsets (Fig. 3F). To explore the role of local DIO2 activity, we incubated endometrial fragments from freshly obtained midluteal biopsies in additive-free medium supplemented with or without T3 for 3 hours (Fig. 3G, fig. S3C and Table S2). T3 treatment did not affect uNK cell marker genes but, as observed in other tissues (*33*), repressed *PLA2G2A* expression in pre-decidual cells, thereby reducing the *PLA2G2A*/*DIO2* ratio by ∼ 50% (Fig. 3H). Further, T3 also inhibited other pre-decidual genes, including *FOXO1*, *SCARA5* and *IL15* (fig. S3D), indicating that DIO2-dependent T3 production confers progesterone resistance in the subluminal implantation niche.

**Fig. 3.**
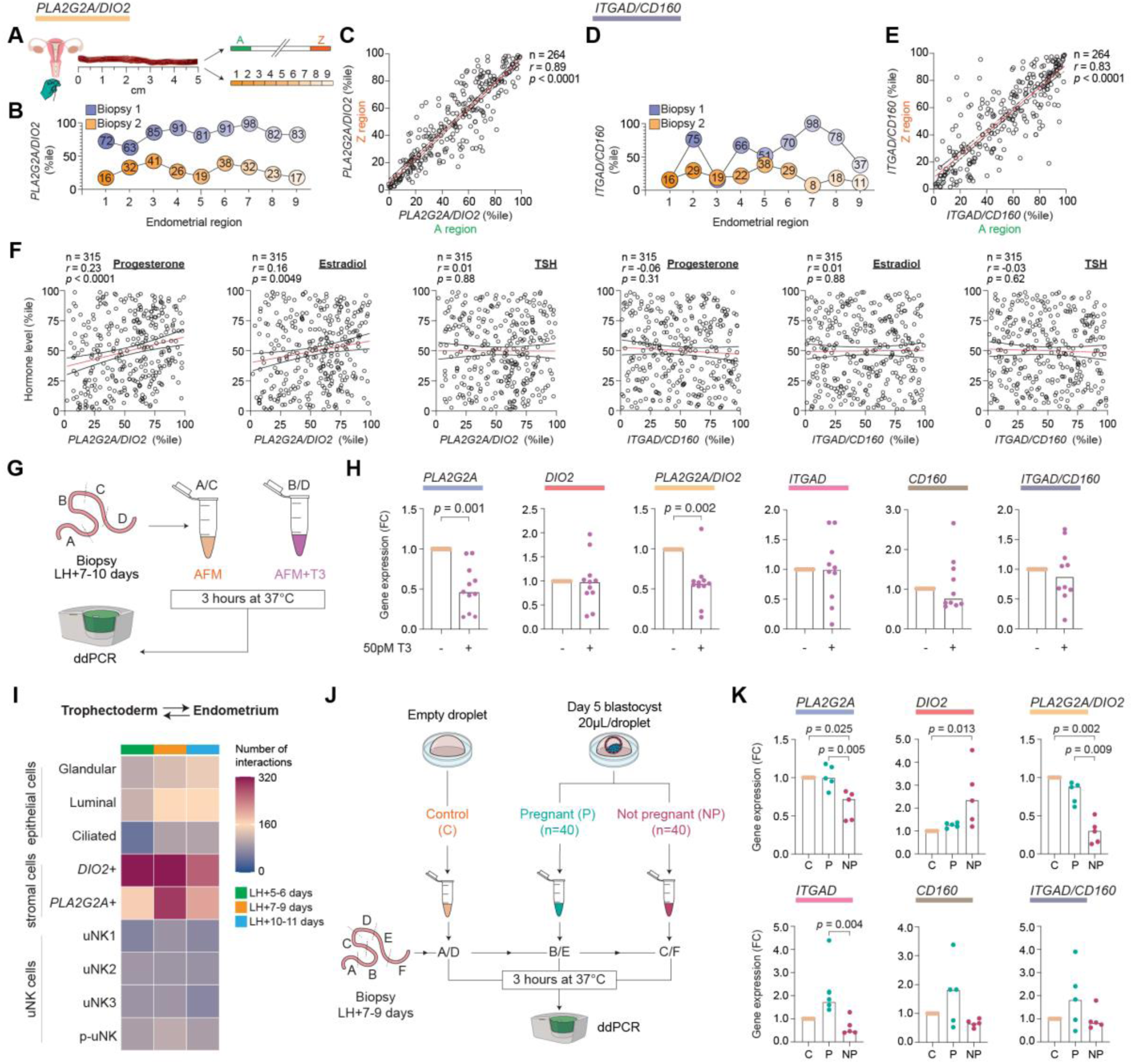
Regulation of stromal and uNK subsets. (**A**) Schematic representation of spatial analysis of normalized *PLA2G2A/DIO2* and *ITGAD/CD160* ratios (%iles) in endometrial biopsies. (**B** and **D**) Representative analysis of two independent samples with normalized gene ratios measured across sequential (∼0.5 cm) regions. (**C** and **E**) Spearman’s correlation of normalized gene ratios at opposite ends of samples (A/Z regions) from 264 subjects. (**F**) Spearman’s correlation between normalized *PLA2G2A/DIO2* or *ITGAD/CD160* ratios (%iles) and circulating progesterone, estradiol and thyroid stimulating hormone (TSH) levels normalized to the day of sampling post-LH surge (%iles). Paired endometrial and peripheral blood samples were obtained from 315 subjects. (**G**) Schematic of experimental design. Freshly isolated endometrial samples were divided, and randomly selected sections incubated for 3 hours in additive-free media (AFM) supplemented or not with 50 pM triiodothyronine (T3) prior to gene expression analysis using digital droplet PCR (ddPCR). (**H**) Bar graphs showing median fold change (FC) in gene expression/gene ratios of 11 and 10 biological repeat experiments for stromal and uNK subsets, respectively. Individual data points are also shown; *p*-values are based on the Wilcoxon matched-pairs signed-rank test. (**I**) Heatmap showing the temporal changes in the number of predicted receptor-ligand interactions between uNK, stromal, or epithelial subsets and trophectoderm cells from day-6/7 human blastocysts. (**J**) Schematic of experimental design. Freshly isolated endometrial samples were divided, and sections incubated for 3 hours in pooled spent medium, diluted 1:1 in AFM, of IVF blastocysts that resulted in a clinical pregnancy (P) or not (NP, not pregnant). Pooled embryo-free droplets served as the control (c) group. (**K**) Bar graphs showing median fold change (FC) in gene expression/gene ratios of five biological repeat experiments. Individual data points are also shown; *p*-values are based on the Friedman with Dunn’s multiple comparison test.

We employed a computational approach to probe how the spatiotemporal changes in endometrial subpopulations could impact embryo implantation. Enumeration of predicted receptor-ligand interactions by CellPhoneDB indicated that the stromal subsets, especially *DIO2*+ cells, play an outsized role in the crosstalk between the endometrium and embryonic trophectoderm cells, the precursors of placental cell lineages (Fig. 3I and Data file S1). Mining of the data for time-sensitive, subset-specific interactions implicated *DIO2*+ cells in decoding trophoblast-derived Wnt signaling at the start of the implantation window and suggested a prominent role for the specialized subluminal ECM in anchoring the implanting embryo (fig. S4A). Finally, we examined if embryonic fitness signals impact on stromal or uNK cell dynamics during the implantation window. Random tissue fragments from midluteal endometrial samples were exposed to pooled spent medium of IVF blastocysts, cultured in individual droplets, which subsequently implanted successfully (n=40) or not (n=40) (Tables, S2 and S3). Unconditioned blastocyst culture medium from ‘empty’ droplets was pooled for control experiments (Fig. 3J). Strikingly, secreted signals emanating from unsuccessful blastocysts enhanced and inhibited *DIO2* and *PLA2G2A* expression, respectively (Fig. 3K). Inhibition of *PLA2G2A* expression was paralleled by repression of *IL15* and other decidual genes (fig. S3E). Spent medium of successful embryos, on the other hand, had no discernible impact on stromal marker genes, but increased *ITGAD* expression (Fig. 3K). Taken together, our observations demonstrate that the balance of *DIO2*+ and *PLA2G2A+* subsets is finely poised during the implantation window, controlled by the opposing actions of circulating progesterone levels and local T3 production. Low-fitness embryos may upend this bistable State in a manner predicted to preclude decidual transformation and promote tissue breakdown. In agreement with previous studies (*15, 34*), expansion of *ITGAD*+ uNK subsets during the implantation window appears regulated by local endometrial factors, such as IL-15 and glycodelin, and likely accelerates upon implantation of a high-fitness embryo.

### The recurrence risk of miscarriage

We reasoned that a stalled decidual response or poor uNK cell expansion in a conception cycle increases the risk of a miscarriage. We first explored this possibility by analyzing endometrial biopsies from 663 subjects selected on the number of previous pregnancy losses (Table S4), a proxy measure of age-independent miscarriage risk (*7, 8*). The frequency of samples with a blunted or stalled decidual response (*PLA2G2A*/*DIO2 <* 25^th^ percentile) increased stepwise with each prior miscarriage, mirrored by a stepwise decrease in samples with a heightened decidual reaction (*PLA2G2A*/*DIO2* > 75^th^ percentile) (Fig. 4A and fig. S5). The frequency of samples with suboptimal uNK cell expansion (*ITGAD*/*CD160* < 25^th^ percentile) increased after three or more losses (Fig. 4B and fig. S5). Women with a stalled decidual response, alone or combined with suboptimal uNK cell expansion, had more prior pregnancy losses (Fig. 4C). There was no association between the number of previous miscarriages and circulating estradiol or progesterone levels (Table S5). However, circulating TSH levels at the higher end of the standard clinical range and increased body mass index (BMI) were associated with higher-order miscarriages (Table S5). Increased BMI was further associated with lower stromal and uNK cell marker gene ratios (Table S6). Next, we analyzed an independent cohort of 261 samples obtained before pregnancy (Tables, S7 and S8). The miscarriage rate in this prospective cohort was 32%, with fetal karyotyping results available for 46 out of 84 miscarriages (Fig. 4D). A stalled pre-pregnancy decidual reaction (*PLA2G2A/DIO2* < 25^th^ percentile) was associated with increased miscarriage risk and decreased likelihood of a livebirth in a future conception cycle (OR: 0.52, 95% CI: 0.29-0.92, *p* = 0.02) (Fig. 4E, left panel). This association was more robust upon omission of confirmed aneuploid losses from the outcome data (OR: 0.42, 95% CI: 0.23-0.77, *p* = 0.005). Exclusion of aneuploid miscarriages also revealed that *ITGAD*/*CD160* ratios > 75^th^ percentile favor live births in a subsequent pregnancy (OR: 2.29, 95% CI:1.05-4.99, *p* = 0.04) (Fig. 4E, right panel).

**Fig. 4.**
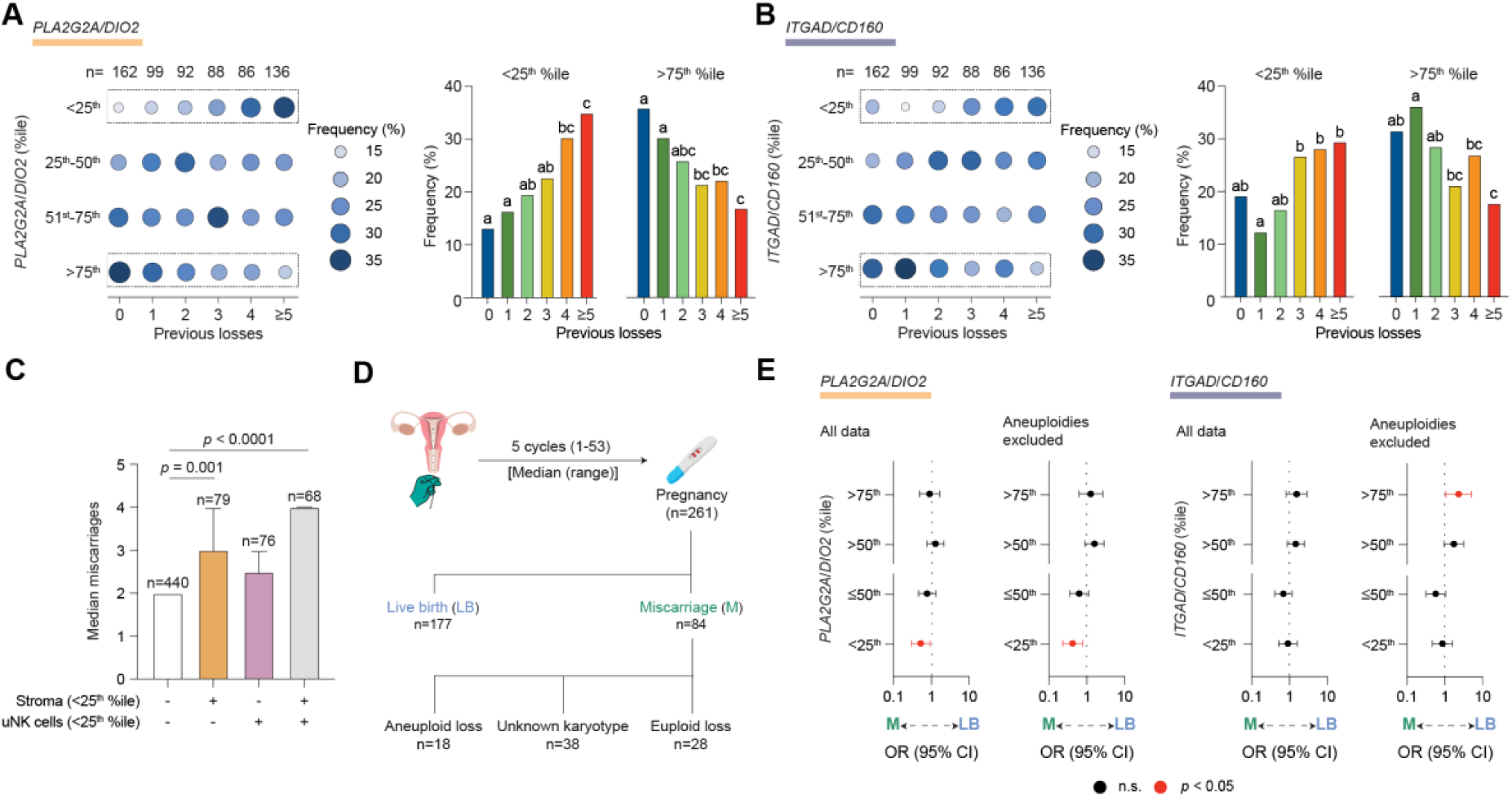
The recurrence risk of miscarriage. (**A** and **B**) Dot plots (left panels) showing the frequency of endometrial samples (%) stratified by the number of previous pregnancy losses for (**A**), *PLA2G2A/DIO2* %iles quartile bins and (**B**), *ITGAD*/*CD160* %iles quartile bins. The number of subjects in each clinical group is also shown. The frequency of samples in the lowest and highest quartile bins is also enumerated in bar graphs (right panels) for (**A**), normalized stromal subset ratios and (**B**), normalized uNK subset ratios. Different letters above the bars indicate significance between groups at *p* < 0.05, Chi-squared test with Bonferroni correction. (**C**) Median number of prior miscarriages in subjects with a stalled decidual reaction (normalized stromal subset ratios < 25^th^ %ile), poor uNK cells expansion (normalized uNK subset ratios < 25^th^ %ile), or both. Statistical analysis is based on one-way ANOVA and Tukey’s multiple comparison test. (**D**) Schematic of the cohort. (**E**) Forest plots showing odds ratios (OR) and 95% confidence intervals (CI) for miscarriage risk and live birth rates following assessment in a non-conception cycle of the decidual reaction (*PLA2G2A/DIO2* %ile) and uNK cell expansion (*ITGAD/CD160* %ile). Subjects were grouped in quartile bins. Data are shown for all subjects (left panels) and upon exclusion of confirmed aneuploid pregnancy losses (right panels); n.s., non-significant.

Our findings imply that cycles characterized by an aberrant endometrial state associated with miscarriage must occur at a higher frequency than expected by chance alone. Analysis of paired endometrial samples obtained in different cycles from 316 women (Fig. 5A, fig. S6 and Table S9) showed that stalling of the decidual reaction (*PLA2G2A*/*DIO2* < 25^th^ percentile) is significantly overrepresented in paired biopsies (Bonferroni-adjusted *p* < 1.0 × 10^-8^) with a recurrence rate of 55% (Fig. 5B). An accelerated decidual response (*PLA2G2A*/*DIO2* > 75^th^ percentile) was also overrepresented (Bonferroni-adjusted *p* < 1.9 × 10^-3^) with a recurrence rate of 36%. By contrast, the level of uNK cell expansion in the first biopsy had no discernible impact on uNK subset dynamics in the second sample (Fig. 5B). Stratification of the data showed that one or two prior miscarriages suffice to disrupt the intercycle dynamics of stromal but not uNK subsets (Fig. 5, C and D). Further, the frequency of a stalled decidual reaction appears to increase in line with the number of prior pregnancy losses (Fig. 5C). Together, these observations highlight that the decidual reaction is intrinsically dynamic, with measurable levels of variation between cycles. Clinically, the incidence of a stalled decidual reaction is a major determinant of the risk of a future miscarriage. Our findings further suggest that prior miscarriages compromise endometrial homeostasis, defined as the ability to maintain the decidual reaction in an optimal range from cycle to cycle.

**Fig. 5.**
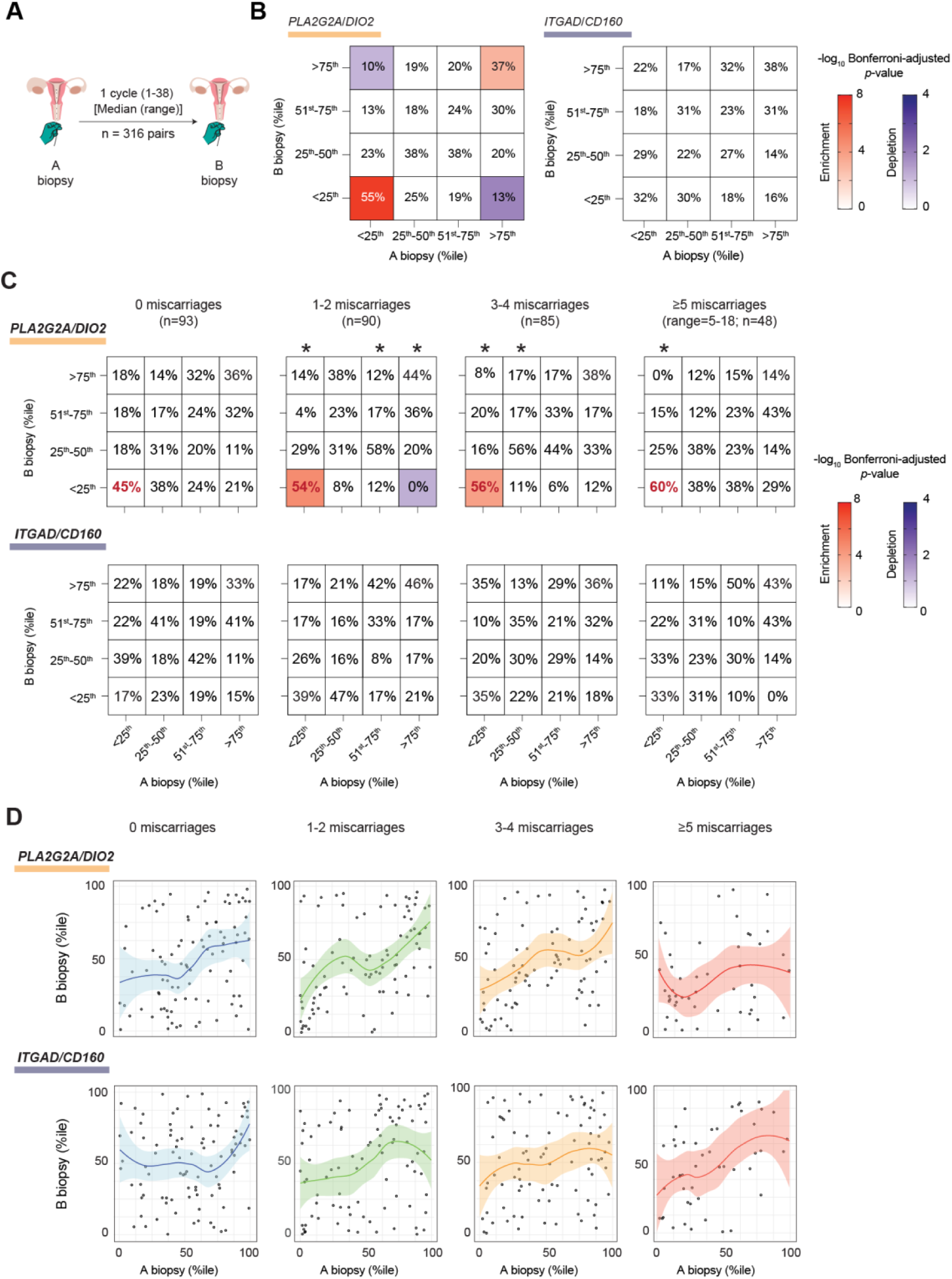
Analysis of intercycle variability. (**A**) Schematic of paired endometrial sample collection. (**B**) Contingency tables of the frequency of samples (%) grouped in quartile bins of normalized stromal and uNK subset ratios (*PLA2G2A/DIO2* %ile and *ITGAD/CD160* %ile, respectively) in paired A and B biopsies. The colored squares in the contingency tables indicate statistical significance (*p* < 0.05) as determined by the Fisher’s exact test for enriched (red key) and depleted (blue key) associations. (**C**) Contingency tables, stratified by the number of prior losses, of normalized stromal and uNK subset ratios (%iles) grouped in quartile bins in paired A and B samples. The color key represents −log10 Bonferroni-adjusted *p*-value as determined by Fisher’s exact test for significantly enriched (red) and depleted (blue) associations. The recurrence rate (%) of a stalled decidual response is indicated in red letters. (*) above a column in the contingency tables indicates non-uniform distribution of the relative frequency of samples in B biopsy quartile bins per A biopsy quartile bin at *p* < 0.05, Chi-squared test. (**D**) Scatter plots with curves of best fit (solid line) and 95% CI (colored areas).

### Impacts of prolonged decidual inflammation

In multiple physiological processes, including tissue repair, the transient release of pro-inflammatory mediators such as IL-6 enhances cellular plasticity by promoting the dedifferentiation of specified cells into stem cells (*35, 36*). However, chronic sterile inflammation causes stem cell depletion and tissue dysfunction (*37*). Miscarriage is linked to senescence-associated inflammation culminating in the destruction of the nascent uteroplacental interface (*38*). To explore how a miscarriage could affect stromal cell responses in subsequent cycles, we model the sequelae of prolonged decidual inflammation in assembloids (Table S2). Following a short pre-assembly culture step, primary stromal and epithelial cells were mixed in a collagen hydrogel and grown into assembloids using a chemically defined expansion medium (ExM) (Fig. 6A and Table S10) (*21*). After eight days of growth (Fig. 6B), the cultures were differentiated for twenty days in minimal decidualization medium (MDM) (Fig. 6C and Table S10). Secretion of IL-6 followed a triphasic pattern with a transient peak around day 4 of decidualization and conspicuous secondary rise from day 10-12 onwards (Fig. 6D), reflecting the absence of immune surveillance of decidual-like senescent cells (*19, 21*). The initial inflammatory decidual response coincided with marked induction in the colony-forming unit (CFU) activity, measuring the relative abundance of clonogenic stem-like cells (*18*). Sustained decidual inflammation, however, abrogated this response (Fig. 6E). We used *SCARA5*/*DIO2* mRNA ratios to monitor progesterone responsiveness, as cultured cells do not express *PLA2G2A*. Prolonged decidual inflammation resulted in progesterone resistance (Fig. 6F) and induction of *MMP10* (Fig. 6G), a matrix metalloproteinase (MMP) expressed by stromal cells at menstruation (*39*). All assembloids contracted and detached spontaneously from the culture plates after 14 days in culture (Fig. 6C). By contrast, assembloids exposed alternatingly every four days to growth and decidualization medium not only maintained their integrity but each ‘cycle’ of decidualization increased CFU activity (Fig. 6, H to I). The data suggest that the decidual reaction enhances tissue plasticity in cycling endometrium. In the context of a clinical miscarriage, however, pathological decidual inflammation sustained over weeks may cause stem cell depletion, progesterone resistance and, plausibly, disruption of the regenerative endometrial-myometrial interface (*2*).

**Fig. 6.**
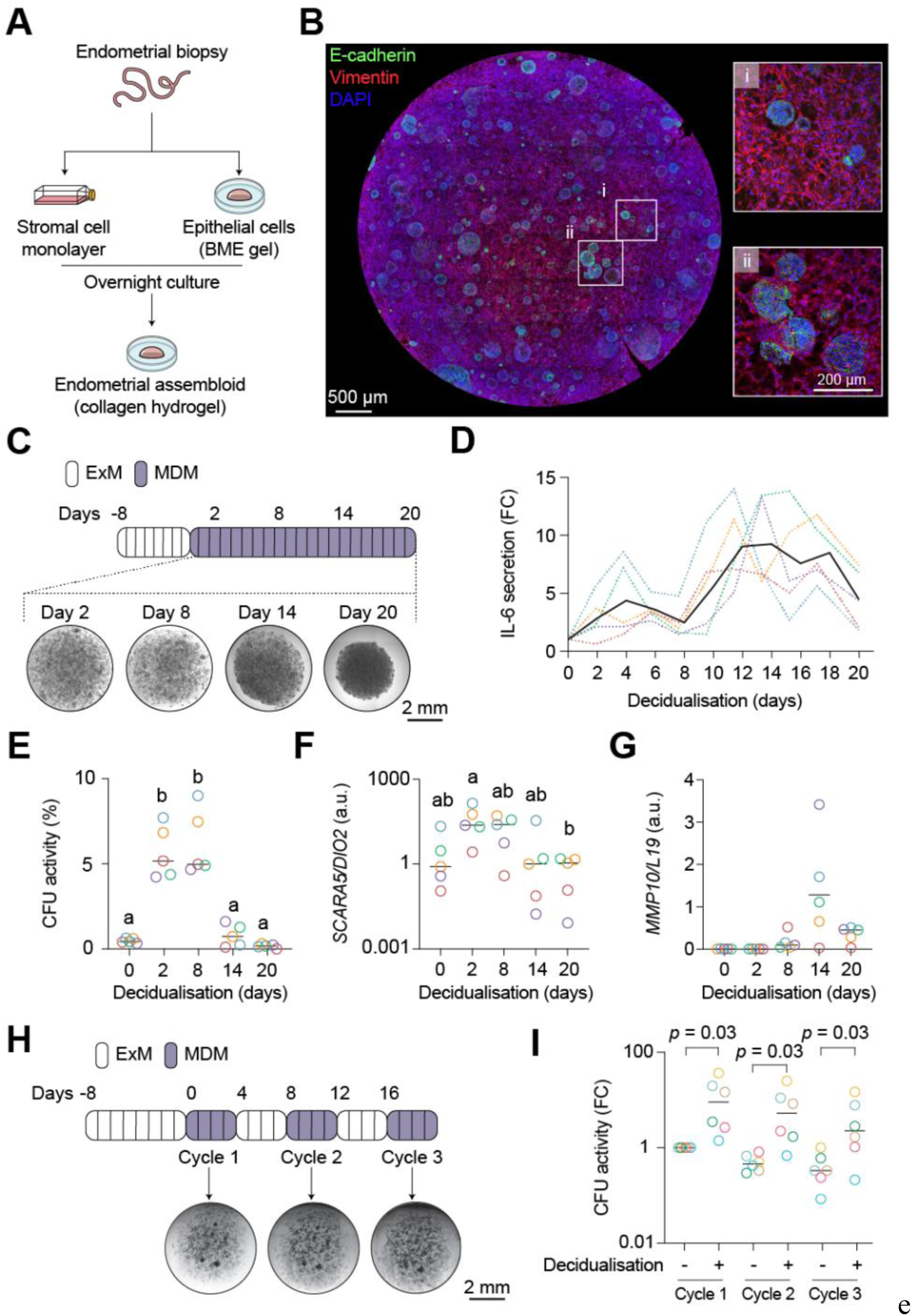
Modelling decidual inflammation in assembloids. (**A**) Schematic workflow for establishing assembloids from primary endometrial stromal and epithelial cells. BME, basement membrane extract. (**B**) Representative confocal image of E-cadherin and vimentin immunofluorescence in an assembloid grown for 8 days in a chemically defined expansion medium (ExM). Nuclei were stained with DAPI. Boxes highlight regions at higher magnification. (**C**) Schematic of experimental design and representative micrographs of assembloids maintained in a minimal decidualization medium (MDM). (**D**) Fold change (FC) in IL-6 secretion upon decidualization of endometrial assembloids. Colored dotted lines represent the FC in secreted IL-6 levels in five biological repeat experiments. The solid black line represents the median FC. (**E**) CFU activity in undifferentiated and decidualized endometrial assembloids. The data show median CFU activity (%) and individual data points of five biological repeat experiments. Different letters above the data points indicate significance at *p* < 0.05, Kruskal-Wallis test with Dunn’s multiple comparisons test. (**F**) Relative change in *SCARA5* and *DIO2* mRNA ratio in five biological repeat experiments. Different letters above the data points indicate significance at *p* < 0.05, Kruskal-Wallis test with Dunn’s multiple comparisons test; a.u., arbitrary units. (**G**) Time-dependent induction of *MMP10* expression. The data show median *MMP10* transcript levels, normalized to *L19*, and individual data points of five biological repeat experiments. (**H**) Schematic of experimental design and representative micrographs of assembloids first grown in ExM for eight days and then alternatingly in MDM and ExM over three cycles. (**I**) Relative change in CFU activity in assembloids across three cycles of decidualization. The data show median FC in CFU activity and individual data points of five biological repeat experiments. Statistical analysis is based on Wilcoxon matched-pairs signed rank test for comparison within each cycle.

## DISCUSSION

The functional integrity of most adult tissues relies on homeostatic circuits that sense and respond to environmental perturbations. Cues that overwhelm the capacity of these circuits trigger sterile inflammation, which serves to restore and recalibrate homeostasis (*40*). The cyclically regenerating endometrium defies the definition of a homeostatic tissue as its core functions - implantation and formation of a stable uteroplacental interface - depend on an endogenous inflammatory reaction, triggered and constrained by progesterone (*2*). By repressing stress-activated signaling pathways, progesterone drives the transcriptional reprogramming of stromal cells into anti-inflammatory decidual cells, which form the placental bed in pregnancy, but also cause menstruation when progesterone levels fall in non-conception cycles (*2, 5*). We identified *PLA2G2A*, an acute-phase inflammatory gene involved in antimicrobial defenses, tissue regeneration and eicosanoid production (*22, 41*), as a sensitive marker gene of progesterone-dependent pre-decidual cells. We also identified acutely stressed source cells as putative drivers of the decidual reaction, although their provenance is uncertain. Spatial mapping revealed that progesterone-resistant *DIO2*+ cells abutting the luminal epithelium constrain the decidual reaction at the start of the implantation window. *DIO2*+ cells express genes involved in EMT and uterine patterning during development. This gene signature implicates Wnt7a, secreted selectively by proliferative phase luminal epithelium (*42*), in imposing positional identity on subluminal stromal cells (*27*). Multiple strands of evidence indicate that *DIO2*+ cells are essential for interstitial implantation, including the overrepresentation of genes predicted to mediate blastocyst-endometrium interactions, the expression of specialized ECM genes, and the inferred motile phenotype. By engaging placental cells, uNK cells are evolutionarily conserved partners of decidual cells (*2, 5, 28*). We observed that the proliferative expansion of uNK cells leads to the crowding of immunotolerant *ITGAD*+ (KIR+) subsets in the implantation niche. By contrast, cytolytic *CD160*+ (KIR−) uNK cells, tasked with curtailing decidual inflammation by pruning acutely stressed and damaged cells (*2, 18*), reside with underlying IL-15-producing *PLA2G2A*+ cells.

Our findings imply that the progressive loss of *DIO2*+ cells and reciprocal expansion of *PLA2G2A*+ cells set the spatiotemporal boundaries of the interstitial implantation niche. In line with this paradigm, an accelerated decidual response is clinically associated with recurrent IVF failure (*43*), whereas delayed implantation beyond the midluteal phase - a predicted consequence of a stalled decidual reaction - markedly increases miscarriage rates (*44*). Although the mechanisms controlling time-sensitive rebalancing of stromal subsets require further investigation, we noted that signals from low-fitness IVF blastocysts, defined by their failure to implant, inhibit the decidual reaction. By contrast, successful embryos produce signals that accelerate the expansion of immunotolerant uNK cells. Thus, by imposing a bistable State on the endometrium, the decidual reaction establishes an ‘implantation checkpoint’ that limits the risk of prolonged maternal investment in low-fitness embryos (*2, 3, 6*).

We report that the age-independent recurrence risk of miscarriage aligns with the frequency of a stalled decidual reaction. We found that suboptimal expansion of immunotolerant uNK cells is associated with higher-order miscarriages, possibly reflecting the impact of comorbidities associated with uNK cell dysfunction, such as obesity (*45*). Notably, strong uNK cell expansion was associated with increased likelihood of a successful future pregnancy. The management of recurrent miscarriage, defined clinically by two or three prior pregnancy losses, focuses mainly on maternal risk factors (*1, 46*). However, the impacts of comorbidities and lifestyle factors on miscarriage rates are modest (*47, 48*), and evidence-based treatments are in most instances not available (*1, 46*). Our findings intimate that prior miscarriages compromise the ability of the endometrium to maintain a calibrated decidual reaction in subsequent cycles, thus enhancing the likelihood of further losses irrespective of embryo ploidy or maternal age. Prolonged decidual inflammation in assembloids leads to progesterone resistance, stem cell depletion, and loss of the structural integrity of the cultures, three intertwined pathological endometrial features linked to miscarriage risk. We previously demonstrated that stem cell depletion in midluteal endometrium corresponds to the number of prior losses (*49*). Imaging studies reported that disruption of the uterine junctional zone, that is, the myometrial portion of the regenerative layer (*2, 50*), is prevalent in recurrent miscarriage (*51, 52*). Inhibitory cytokines emanating from tertiary lymphoid structures in the regenerative layer ensure that endometrial cells become more hormone-responsive with increasing distance from the endometrial-myometrial interface (*2*). Hence, progesterone resistance, culminating in a poor decidual reaction, may partly reflect the loss of spatial patterning caused by disruption of the regenerative layer. In line with this conjecture, a recent prospective study reported that sonographic evidence of junctional zone disruption does not affect implantation rates but significantly increases miscarriage risk independently of embryo quality (*52*).

Our study has limitations. We evidenced the biological plausibility that a miscarriage increases the risk of further pregnancy losses directly by disrupting the mechanisms that govern intercycle homeostasis. These mechanisms, however, remain poorly understood and causal evidence requires carefully designed longitudinal studies and interventional clinical trials. Further, *miscarriage* is an umbrella term that encompasses different clinical presentations (*1, 53*). Stalling of the decidual reaction is anticipated to cause the breakdown of the nascent uteroplacental interface and bleeding in early gestation. Thus, the relevance of our findings to other recurrent clinical presentations, such as missed (silent) miscarriages, is unclear. Finally, our study focuses on stromal and uNK cell dynamics in midluteal endometrium; the contributions of other cellular constituents, including epithelial, vascular and lymphoid populations and the endometrial microbiome (*54*), to miscarriage risk warrant further exploration.

In summary, the incidence of chromosomal errors in oocytes and embryos accounts for the rise in miscarriage rates with advancing maternal age (*10, 11*). Here, we report that the frequency of menstrual cycles resulting in a suboptimal decidual reaction determines the age-independent recurrence risk. We posit that the likelihood of these two independent events - implantation of an aneuploid embryo and stalling of the decidual reaction - occurring in a given conception cycle explains miscarriage risk in line with epidemiological observations and accounts for the generally high cumulative live birth rates following multiple pregnancy losses (*3, 8, 55*). This disease paradigm also explains why treatments initiated in pregnancy that target subclinical maternal comorbidities are generally ineffective (*1, 46*). Our findings provide the foundations for the development of endometrial tests to identify women at risk of miscarriage and evaluate pre-pregnancy therapeutics. When combined with other emerging miscarriage diagnostics, such as routine assessment of fetal ploidy status using circulating cell-free fetal DNA-based testing (*56*), novel strategies that improve pregnancy outcomes for the many women and their partners affected by miscarriage are bound to emerge.

## MATERIALS AND METHODS

### Study design

The primary objective of this study was to elucidate the risk of miscarriage imparted by a suboptimal decidual reaction. To achieve this goal, we investigated the cellular dynamics across the implantation window, established methods to measure the decidual reaction and expansion of immunotolerant uNK cells and conducted mechanistic investigations using primary cultures, *ex vivo* tissue samples and assembloids. We delineated the impacts of spatial and intercycle variation, hormone levels, circulating immune cells and embryo signals on the dynamics of stromal and uNK subsets in midluteal endometrium and assessed the decidual reaction and uNK cell expansion in defined patient cohorts.

This study was conducted at the Implantation Research Clinic, a dedicated experimental reproductive medicine clinic at University Hospitals Coventry and Warwickshire (UHCW) National Health Service (NHS) Trust, Coventry, UK. Patients attending the clinic for endometrial assessment presented with a history of one or more miscarriages and/or fertility issues, including unexplained, male-factor and tubal infertility. We defined *miscarriage* as the spontaneous loss of pregnancy up to 24 weeks of gestation, excluding pregnancies of unknown location, ectopic or molar pregnancies, and terminations for any reason. We adopted a broad definition of miscarriage encompassing all spontaneous pregnancy losses without a requirement for ultrasound confirmation in line with the recommendations from an international consensus development study on core outcome sets for miscarriage studies (*57*). Surplus endometrial and peripheral blood samples obtained for diagnostic purposes were processed for research. The use of surplus samples for research was approved by the Arden Tissue Bank at UHCW NHS Trust (NHS Research Ethics Committee approval: 18/WA/0356). All samples were obtained following written informed consent and in accordance with The Declaration of Helsinki (2000) guidelines. Patients in the prospective outcome cohort (Table S7) were enrolled in Tommy’s Net (ISRCTN registry: ISRCTN17732518), a dedicated electronic database for miscarriage research (*58*). A total of 1555 endometrial biopsies from 1308 women were processed for this study. In addition, 322 blood samples were obtained from 301 women obtained in ovulatory cycles, defined by serum progesterone levels > 10 nmol/L (*59*). All study participants had circulating TSH levels within the clinical reference range (0.55-4.78 mIU/L). A minimal set of anonymized demographic data was collected for each endometrial sample (see Tables S1 to S9). The use of spent blastocyst culture medium was approved by the Institutional Ethical Committee of the UZ Brussel, Belgium (BUN1432021000527). Anonymized samples were pooled based on clinical outcomes (Table S3). Spent medium of IVF blastocysts (n=40) that resulted in negative pregnancy test 14 days following single embryo transfer were allocated to the ‘not pregnant’ group. The ‘pregnant group’ comprised pooled culture droplets of single embryos (n=40) that resulted in a positive pregnancy test and ultrasound evidence of a viable clinical pregnancy at seven weeks of gestation. Data were acquired in a blinded manner but unblinded during data analysis.

### Statistical analysis and visualizations

Statistical analyses were performed using R (v 4.4.0) and GraphPad Prism (v 10.2). Relative expression of stromal cell (*PLA2G2A*, *DIO2*) and uNK cell (*ITGAD*, *CD160*) marker genes was determined in 791 biopsies by log-transforming droplet digital polymerase chain reaction (ddPCR) copy numbers. The normalized expression per LH+day was fitted to a normal distribution using the ‘fitdistr’ and ‘fitdist’ functions from the MASS (*60*) and fitdistrplus (*61*) packages. Ratios were fitted against a gamma or a log-norm distribution depending on Cramér-von Mises criterion per LH+day. Quantiles were log-transformed. Chi-squared test of independence was used to test for significant enrichment or depletion of *PLA2G2A*/*DIO2* and *ITGAD*/*CD160* percentile bins across paired endometrial samples. For each bin, a 2×2 grid was tested using a one-vs.-all approach. Fisher’s exact test with Bonferroni correction was applied on stratified miscarriage data with *p* < 0.05 considered significant. Chi-squared goodness of fit test was used to test uniformity of data in B biopsies for each A biopsy percentile bin. Curves of best fit for paired biopsies stratified by number of miscarriages were produced using ggplot2’s geom_smooth with default parameters. Multiple linear regression analysis was conducted using DATAtab (DATAtab, Graz, Austria). The model was estimated using the ordinary least squares (OLS) method. The DATAtab output provided regression coefficients (β), standard errors, *t*-values and *p*-values. These were used to assess model significance and explanatory power, with statistical significance determined at a < 0.05. All other statistical analyses encompassed Shapiro-Wilk test for normality followed, as appropriate, by paired *t*-test, Wilcoxon Signed-Rank test for paired data, one-way ANOVA, Kruskal-Wallis test with Dunn’s test for multiple comparisons, or one-way ANOVA or Friedman test for data with three or more groups. Visualizations were performed using ggplot2 (v 3.5.1) in R or on GraphPad Prism.

## List of Supplementary Materials

Materials and Methods

Fig. S1 to S6

Tables S1 to S15

Date file S1: Bulk and single-cell RNA-seq analysis and receptor-ligand predictions.

Date file S2: Source files

## ACKNOWLEDGEMENTS

We thank the patients attending the Implantation Research Clinic at University Hospitals Coventry and Warwickshire (UHCW) National Health Service (NHS) Trust for contributing endometrial and blood samples for research. We also thank the staff of the Biomedical Research Unit in Reproductive Health, Tommy’s National Miscarriage Research Centre, Arden Tissue Bank, Coventry and Warwickshire Pathology Services and Research and Development Department at UHCW NHS Trust. We are indebted to Amelia Hawkes, Joshua Odendaal, Lauren Lacey, Lauren J. Ewington, Shreeya Tewari, Ioannis Pavlidis, Majd Alkhoury, and Natalie Morris for assisting with sample collection and processing. We thank Komal Makwana, Niky Moolchandani Adwani and Joe Thornton for technical support, and Simon Spencer for statistical advice.

## Author contributions

Conceptualization: J.J.B.; Funding acquisition: C.-S.K., S.O., J.J.B; Investigation: J.M., C.-S.K., M.T.N., M.T., P.V., P.J.B, D.B.D., M.V., E.S.L., T.M.R., Methodology: J.M., C.-S.K., M.T.N, M.T., P.V., H.Y., S.O., B.K.T., P.R.B., S.Q., A.R., H.V.d.V., T.M.R.; Visualization: J.M., C.-S.K., M.T.N., P.V., Supervision: S.O., S.Q., A.R., H.V.d.V, J.J.B.; Writing – original draft: J.J.B; Writing – review & editing: J.M., C.-S.K., B.K.T., P.R.B., E.S.L., H.V.d.V., T.M.R.

## Data and materials availability

All data included in the study are provided in Data files S1 and S2. The bulk RNA-seq data and scRNA-seq data generated in this study were deposited in the NCBI Gene Expression Omnibus repository (https://www.ncbi.nlm.nih.gov); accession numbers GSE245963 (bulk RNA-seq data) and GSE247962 (scRNA-seq data).

## Materials and Methods

### Endometrial sample collection

Anonymized endometrial samples were obtained during the luteal phase of ovulatory menstrual cycles and timed relative to the pre-ovulatory LH surge, as determined by commercially available ovulation test kits. Overt uterine pathology was excluded by vaginal ultrasonography scan prior to the biopsy. The thickness of the endometrium was measured at the maximum distance between each myometrial/endometrial interface perpendicular to the sagittal axis of the uterus. Endometrial biopsies were obtained using a Wallach Endocell™ or CerviX™ endometrial sampler.

### Blastocyst conditioned media

Spent culture medium was collected following *in vitro* culture of human embryos created either by *in vitro* fertilization (IVF) or intracytoplasmic sperm injection (ICSI) following controlled ovarian stimulation and oocyte retrieval (*62, 63*). Human embryos were cultured for 5 days in individual 25 μL droplets of ORIGIO® Sequential Series™ medium, overlaid in OVOIL™ (Vitrolife, 10029) at 37°C, 6% CO_2_, 5% O_2_ in a G185 flatbed incubator (CooperSurgical, Denmark). Embryos were first cultured for 3 days in 25 μL droplets of ORIGIO® cleavage-stage medium under oil and then cultured in 25 μL droplets of ORIGIO® blastocyst medium under oil. Single day-5 embryos, selected based on morphological criteria (*64*) (Table S3), were transferred into the uterine cavity. Spent culture medium (22 μL) was collected following embryo transfer using a sterile MµltiFlex pipette tip (Sorenson BioScience, 13810), transferred to a nuclease-free cryovial (Nalgene, USA), snap frozen in liquid nitrogen and stored at −80°C.

### RNA extraction and cDNA synthesis

RNA extraction from RNAlater-preserved tissues was performed using the RNeasy Plus Universal Mini Kit according to the manufacturer’s protocol (Qiagen, 73404). RNA concentrations were determined using a Nanodrop ND-1000 spectrophotometer, and equivalent RNA quantities reversed transcribed into cDNA using the QuantiTect Reverse Transcription Kit (Qiagen, 205314).

### RT-qPCR

Amplification of target genes by RT-qPCR was performed on a QuantStudio 5 Real-Time PCR system (Applied Biosystems, Paisley, UK). cDNA transcribed from RNA (1 µg) was diluted to a concentration of 10 ng/µL. For the qPCR reaction, 1 µL of cDNA was combined with 9 µL of reaction mixture, which included QuantiNova SYBR Green Master Mix (Qiagen, 208057), a low concentration of ROX Dye solution, and 300 nM each of forward and reverse primers. The primer sequences are provided in Table S11.

### Droplet digital polymerase chain reaction (ddPCR)

ddPCR was performed on the QX200 AutoDG Droplet Digital PCR System (Bio-Rad). Briefly, 10 ng of cDNA was added to a 19 μL reaction mixture containing 1 × supermix for probes, 900 nM target primers, and nuclease-free water. Target probes are provided in Table S12. Droplet generation was accomplished using the automated droplet generator (Bio-Rad). The droplet emulsion was subjected to thermal cycling under the following conditions: one cycle of enzyme activation (95°C, 30 sec), 40 cycles of denaturation (94°C, 30 sec), annealing/extension (55°C, 1 min) and one cycle of enzyme deactivation (98°C, 10 min). Post-cycling, PCR amplification within the droplets was assessed using the QX200 Droplet Reader (Bio-Rad). Gene expression threshold between positive and negative drops were determined using QuantaSoft software (version 2.1).

### Bulk mRNA library preparation and sequencing

RNA-seq libraries were mapped to the hg38 human genome assembly using STAR v2.5.3a (*65*) using default settings and gencode.v38 as annotation file (Ensembl). Differential gene expression analysis was performed with DESeq2 v1.34.0 in R (*66*). Principle component analysis (PCA) was performed with prcomp function following transformation of raw counts using the rlog function.

### Processing endometrial samples for scRNA-seq

Endometrial biopsies in additive-free DMEM/F12 media were processed within 15 min of collection. Samples were mechanically minced and enzymatically digested with DNase I (100 µg/mL), Collagenase Type Ia (500 µg/mL) and IV (10 µg/mL) for 90 min at 37°C, with shaking every 15 min. Undigested tissue was removed through a 40 μm cell strainer. To eliminate red blood cells, the cell suspension was incubated with red blood cell lysis buffer (Invitrogen, 00-4333-57) for 5 min at room temperature. The reaction was terminated by adding 30 mL PBS and centrifuged at 400 × *g* for 5 min. Cells were then processed according to Demonstrated Protocol CG00054 (Revision B, 10X Genomics) with some modifications. Briefly, cells were centrifuged at 300 × *g* for 5 min and resuspended in 0.04% BSA in PBS. This was repeated twice more before cells were resuspended in 500 μL PBS with 0.04% BSA and counted. Cells were adjusted to 700 cells/μL.

### Single-cell capture and library preparation

Freshly digested endometrial cells were encapsulated into emulsion droplets using the 10X Genomics Chromium Controller and NextGEM Single Cell 3’ Reagent Kit (v3.1), according to the user guide CG000315, Rev B, targeting 8000-10000 cells per run. Libraries were constructed as per manufacturer’s instructions. Quality control and library quantitation were performed using the Agilent Bioanalyzer and KAPA Library Quantification Kit (KAPA Biosystems, KK4824). Sequencing was performed on a NextSeq 500, using Illumina NextSeq500/550 High Output Kits, v2.5 (150 cycles) with read lengths as described by 10X Genomics (Read 1: 28, i7 Index: 10, i5 Index: 10, Read 2: 90), targeting 20,000 read pairs per cell.

### Single-cell data pre-processing and quality control

Raw sequencing data was processed using CellRanger v6.1 pipelines `cellranger mkfastq` and `cellranger count` (10X Genomics). SoupX (*67*) was used to remove ambient RNA contamination, and count matrices were processed using Seurat v4.4 (*68*). Cells were excluded from the analysis if they had less than 200 detected genes or more than 25 % mitochondrial gene expression. DoubletFinder (*69*) (version 2.0.4) was used to predict and remove cell doublets. Integration was performed across patient samples using the reciprocal PCA method. A total of 64,644 high-quality endometrial cells were retained for further bioinformatic analysis.

### Single-cell data integration, dimensionality reduction and cell annotation

Global scaling normalization was applied through the NormalizeData function. Highly variable genes were identified through FindVariableGenes function. PCA was performed on the integrated dataset. Top 20 PCs were selected for downstream analysis with the Elbowplot function. The main cell clusters were identified with the Seurat FindClusters function with the resolution set at 0.5. The endometrial cells were clustered and visualized by uniform manifold approximation and projection (UMAP) plots. scCustomize package (*70*) was used to visualize gene expression. Cell type identification was performed by manual inspection of marker genes and interpretation of these based on previous studies. Subsequently, stromal cells and uNK cells were further subclustered to detect heterogeneity. The differential gene expression analysis of identified cell clusters was executed using the Wilcoxon rank-sum test within the Seurat’s FindAllMarkers command. We used the GSEA (*71*) with hallmark gene sets to identify biological pathways enriched in endometrial stromal subsets (Data file S1).

### Receptor-ligand interactions

CellPhoneDB v4 (*72*) was used to predict enriched receptor-ligand interactions involved in the crosstalk between human trophectoderm from day-6/7 blastocysts and endometrial subsets. Endometrial subpopulations were identified based on thresholding (>0) of marker gene counts. scRNA-seq data from human embryos (E-MTAB-3929) (*73*) were normalized using Seurat’s SCTransform and then clustered using FindClusters at resolution 0.5. Clusters were identified as trophectoderm, epiblast or hypoblast through VlnPlot using relevant biomarkers. The datasets were merged, and counts were globally normalized using SCTransform. Annotated temporal interactions were curated manually. The receptor-ligand interactions are listed in Data file S1.

### Single-cell RNA velocity

RNA velocity analysis, which calculates the ratio of unspliced to spliced RNA for each gene, was performed to identify the direction of differentiation (*74, 75*). Fastq files were first processed using loompy Python package v3.0.7 (fromfq using gencode. v31.600. Kallisto v0.46.2). To estimate RNA velocity from scRNA-seq data, we utilized the scVelo package (*75*) v0.2.4 in Python v3.9.13. A velocity vector field was constructed for all cells in the dataset, where each cell was assigned a vector direction and magnitude representing its predicted future gene expression state. Source cells were identified as 100 cells closest to the origin of the velocity vectors. Velocities were plotted into UMAP space.

### Single-cell analysis of transposable element (TE)-derived transcripts

Locus-specific quantification of TE-derived transcripts in the transcriptomic data was performed with SoloTE (*76*). The resulting count matrix was used for single-cell analysis using Seurat.

### Multiplexed single molecule in situ hybridization (smISH)

Endometrial biopsies were fixed overnight in 10% neutral buffered formalin. Tissue processing through graded alcohol and embedding in Surgipath Formula ‘R’ paraffin was automated using the Shandon Excelsior ES Tissue Processor, and a Tissue-Tek TEC embedder. Tissues were sectioned at 5 μM and mounted on SuperFrost Plus slides (Epredia, J1800AMNZ). Staining was carried out using the RNAscope Multiplex Fluorescent Reagent Kit v2 Assay (Advanced Cell Diagnostics, Bio-Techne). The probes are listed in Table S13.

### Visium spatial transcriptomics

Visium slides and library preparation / Read mapping Spatial transcriptomics were carried out using the 10x Genomics Visium platform. Spaceranger software (version 1.3.0, 10x Genomics) was used to align and obtain raw counts from each of the spots on the Visium spatial transcriptomics slides against the GRCh38 human genome reference data (refdata-gex-GRCh38-2020-A). The spatial transcriptomics raw gene expression matrix, together with spatial location of spots and tissue hematoxylin and eosin (H&E) images, were used to create a Seurat object with a Load10X_spatial function. After normalization by SCTransform, we performed principal component analysis and reduced the dimensions to the top 20 principal components. Marker gene detection and differential gene expression were carried out using the FindAllMarkers function in Seurat. Genes that varied by location were identified using the FindSpatiallyVariableFeatures function using default settings. The DNA repair gene signature was visualized by summing the count data of the 150 genes in the GSEA HALLMARK_DNA_REPAIR set(*71*).

### Serum hormone levels

Blood was collected immediately prior to biopsy collection in serum-separating tubes. Quantitation of hormonal levels in serum was based on acridinium ester chemiluminescent technology using the Atellica IM Analyser system (Siemens Healthcare GmbH, Erlangen, Germany). TSH (Atellica IM TSH3 UL) was quantified through direct analysis utilizing FITC-labelled anti-TSH capture mouse monoclonal antibody, this was bound to the solid phase via an anti-FITC monoclonal antibody. Introduction of an anti-mouse monoclonal antibody labelled with acridinium initiated the reaction and a direct relationship was observed between the amount of TSH in the sample and relative light units detected. Progesterone (Atellica IM PRGE assay) and estradiol (Atellica IM E2 assay) were measured using a competitive immunoassay format. The target analyte was first bound to an acridinium-ester-labelled monoclonal anti-hormone antibody. A derivative of the target hormone coupled to the capture solid phase was then introduced, which competes and binds any remaining antibody. Following a wash, the addition of acid and base to initiate the reaction resulted in an inverse relationship between the amount of hormone in the sample and relative light units detected. Samples were processed in a fully UKAS accredited medical diagnostic laboratory working to the international ISO 15189:2012 standard.

### Immunohistochemistry

Formalin-fixed, paraffin-embedded human endometrial tissue sections (3 µm thick) were deparaffinized three times with xylene, 5 min each, and rehydrated using 100% isopropanol (2×) and 70% isopropanol (1×), each for 5 min. Antigen retrieval was performed in citrate buffer (10mM citrate, 0.05% Tween 20, pH 6). After blocking with 2% BSA in TBST, sections were incubated with indicated primary antibodies overnight at 4°C. Subsequent washing with TBST was followed by incubation with Alexa-fluor 594 anti-mouse or Alexa-fluor 488 anti-rabbit secondary antibodies for 2 hours at room temperature. After three TBST washes, the sections were treated with Vector TrueVIEW auto-fluorescent quenching solution (Vector Laboratories, SP-8400). The slides were washed with PBS and cover slipped using the ProLong™ Gold Antifade Mountant with DAPI DNA Stain. Imaging was conducted using a BioTek Cytation C10 confocal imaging reader (Agilent Technologies) and analyzed using Gen5 image analysis software version 3.12. ImageJ/Fiji software was used for editing purposes (*77, 78*). The antibodies are listed in Table S14.

### Flow cytometry and sorting of primary uNK cells

Following tissue digestion, endometrial single-cell suspension was incubated with red blood cell lysis buffer (Invitrogen, 00-4333-57) for 5 min at room temperature. The reaction was terminated by adding 30 mL PBS and centrifuged at 400 × *g* for 5 min. The single-cell suspension was washed twice with PBS (300 × *g* for 5 min) and incubated with fluorescent-conjugated antibodies in a wash buffer (0.5% BSA in PBS with 2.5 mM EDTA, 4°C). Live and dead cells were discriminated using Fixable Viability Stain 660 (1:1000) in PBS. Cell phenotyping and sorting were conducted on a Becton Dickinson (BD) FACS Melody with 3 lasers (488 nm, 561 nm, and 640 nm) and 8 detectors. Sorting utilised BD FACS Chorus software (version 3.0). Analyses were performed using FlowJo software (version 10.9.0). Antibodies are listed in Table S15.

### Peripheral blood lymphocyte immunophenotyping

Whole blood drawn into BD EDTA Vacutainers and BD Vacutainer® CPT™ Mononuclear Cell Preparation Tubes with Sodium Heparin (BD Biosciences, 362753) was used for T, B, and NK cell (TBNK) assays (BD Biosciences, 644611). A lyse/no-wash method was employed by adding 20 μL of BD Multitest™ 6-color TBNK Reagent to BD Trucount™ tubes (BD Biosciences) and mixing it with 50 μL of anticoagulated whole blood. The tubes were gently vortexed and incubated at room temperature for 15 min. Subsequently, 450 μL of Lysing solution (BD Biosciences) was added, followed by another 15 min incubation at room temperature. The samples were then analyzed using the BD FACSCanto™ II cytometer, and population numbers were calculated with BD FACSDiva™ Software (version 9.2). Leukocytes were gated based on CD45 positivity and side scatter. Within this gate, the relative abundance of CD56+/CD16+ NK cells and CD3+ T cells was quantified.

### Endometrial explant culture

Fresh endometrial tissue samples divided into pieces within 1-2 min of sample collection were added to 1.5 ml Eppendorf tubes containing additive-free DMEM/F12 medium with or without T3 (50 pM) or blastocyst conditioned medium (1:1 dilution with additive-free media). Samples were maintained at 37 °C in 5% CO2 on a rocking platform with a frequency of 0.1 Hz. After 3 hours, tissues were immersed in RNAlater for RNA extraction, cDNA synthesis, and ddPCR analysis.

### Isolation of primary stromal and epithelial cells

Endometrial biopsies were chopped using scalpels for 5 min and enzymatically digested in 5 mL of pre-warmed additive-free DMEM/F12 medium containing 500 µg/mL Collagenase Type Ia (Merck Life Science, C9891) and 100 µg/mL DNase I (Lorne Laboratories LTD, LS002060) for 1 hour at 37 °C. Cell suspension was then filtered through 40 µm cell strainer to separate epithelial cells. The flowthrough contained stromal cells, and the strainers were inverted and back-washed to collect epithelial cells. Cells were centrifuged (300 × *g*) for 5 min and then resuspend in 10% DMEM/F12 growth medium (composition: DMEM/F12 with phenol red supplemented with 10% dextran-coated charcoal-stripped foetal bovine serum (DCC-FBS), Antibiotic-Antimycotic mix (Gibco, 15240062), 2 mM L-glutamine (Gibco, 25030081), 1 nM β-estradiol (Merck Life Science, E2758) and 2 μg/mL recombinant human insulin (Merck Life Science, 91077C). Cells were routinely cultured at 37 °C in a 5% CO_2_, humidified environment.

### Decidualization of primary stromal cells and uNK cell co-cultures

Non-passaged endometrial stromal cells grown to confluency were then incubated overnight in 2% DMEM/F12 reduced-serum media (composition: DMEM/F12 medium without phenol red supplemented with 2% DCC-FBS, Antibiotic-Antimycotic mix and 2 mM L-glutamine) to synchronize the cell cycle. For decidualization, cells were treated with 0.5 mM 8-bromo-cAMP (Merck Life Science, B7880) and 1 μM medroxyprogesterone acetate (MPA) (Merck Life Science, M1629) in 2% DMEM/F12. For stromal-uNK cell co-cultures, KIR^+/-^ uNK cell subsets were isolated by FACS and added (50,000 events/well) directly to primary stromal cell cultures first decidualized for 6 days in 96-well plates. The co-cultures further treated with 0.5 mM 8-bromo-cAMP and 1 μM MPA for two additional days. Protein lysates were then harvested for quantitation of SA-β-Gal activity.

### SA-β-Gal activity quantitation

SA-β-Gal activity was measured using the 96-well Quantitative Cellular Senescence Assay kit (Cell Biolabs Inc, San Diego, U.S.A). Endometrial stromal cells were rinsed with ice-cold PBS and lysed in 50 μL of ice-cold lysis buffer containing protease inhibitors (cOmplete Protease Inhibitor Cocktail, Roche). Subsequently, 25 µL of 2 × assay buffer was added to 25 µL of cell lysates transferred into black-walled, black-bottomed 96-well plates. The plates were sealed and incubated for 1 hour at 37°C. The reaction was terminated by the addition of 200 μL stop solution. Fluorescent intensity, measured by arbitrary fluorescent intensity units (F.I.U), was determined using a PHERAstar FS plate reader (BMG Labtech) at 360/465 nm.

### Endometrial assembloids

Assembloids were established as described previously(*21*) with some important modifications. Following the isolation of primary stromal and epithelial cells from endometrial biopsies, stromal cells were cultured as described above. Freshly isolated endometrial gland fragments were resuspended in 500 µL phenol red-free DMEM/F12 medium in a microcentrifuge tube and centrifuged at 600 × *g* for 5 min. The medium was aspirated and ice-cold, growth factor-reduced Cultrex RGF Basement Membrane Extract, Type 2 (BIO-TECHNE) was added at a ratio of 1:20 (cell pellet: Cultrex). Samples were aliquoted in 20 µL droplets to a 48-well plate, one drop per well, and allowed to cure for 30 min and then overlaid with 200 μL of expansion medium (ExM). Stromal cells and gland organoids were digested, as described previously(*21*), into single cell suspension 24 hours after establishment and mixed at a ratio of 1:1 (v/v) and ice-cold PureCol EZ Gel (Sigma-Aldrich) added at a ratio of 1:20 (cell pellet: hydrogel). The suspension was aliquoted in 20 μL droplets into a 48-well plate (1 droplet per well). Following hydrogel polymerization at 37 °C and 5% CO_2_ for 60 min, droplets were overlaid with 200 μL of ExM with 10 nM β-estradiol. Medium was refreshed every 48 hours for 8 days. To induce decidualization, ExM was replaced with minimal decidualization medium (MDM) containing 1 nM β-estradiol, 0.5 mM 8-bromo-cAMP and 1 μM MPA. The composition of ExM and MDM is tabulated in Table S10.

### Imaging of endometrial assembloids

For immunofluorescent microscopy, assembloids were fixed in 4% paraformaldehyde in PBS for 15 min at room temperature to crosslink proteins and preserve cellular integrity. The procedure was carried out in 1.5 mL Eppendorf tubes. After fixation, assembloids were washed 3 times with PBS-0.05% Tween 20 for 5 min and permeabilized in PBS-0.1% TritonX for 1 hour at room temperature. Assembloids were incubated in primary antibody (Table S14) diluted in antibody/blocking buffer (10%FBS, 2%BSA, 0.05%Tween 20 in PBS) overnight at 4 °C. Assembloids were then washed in PBS-0.05% Tween 20 for 3 × 5 min before incubation with the secondary antibody for 1 hour in the dark at room temperature. Following washing with PBS-0.05% Tween 20 (3 × 5 min), assembloids were incubated in Hoechst 33342 for 20 min, treated with autofluorescence quenching solution and imaged on ibidi μ-slide 8-well chamber slides (ibidi GmbH, 80826). Samples were kept in clearing solution for 3-5 days. Immunolabelled assembloids were imaged using a confocal microscope (LSM800 equipped with Airyscan, ZEISS) with z-stack scans to reconstruct the 3D structure. Z-stacks were processed with Zen-Blue software (ZEISS). The EVOS M7000 Imaging System (Thermo Fisher Scientific) was used for brightfield and phase contrast microscopy. ImageJ/Fiji software was used for editing purposes (*77, 78*).

### Colony forming unit (CFU) assay

For stromal cell isolation, assembloids were washed with PBS before cell recovery by scraping. Samples transferred into microcentrifuge tubes were centrifuged at 600 × g for 6 min, resuspended in 500 µL of 500 µg/mL Collagenase I and 100 μg/mL DNase I diluted in additive-free DMEM/F12, and incubated at 37°C for 10 min with regular manual shaking to release the cells from the hydrogel. Samples were then centrifuged (600 × g, 6 min, room temperature), the pellets resuspended in 5 mL phenol red-free DMEM/F12 growth medium, and the mixtures passed through a 40 μM cell strainer to isolate stromal cells. Stromal cells were pelleted by centrifugation at 300 × *g* for 5 min and resuspended in 1 mL growth medium. Viable isolated stromal cells were counted in trypan blue on a Neubauer Improved hemocytometer, seeded at clonal density (104 cells/cm^2^) onto 6-well plates coated with fibronectin (2 μg/cm^2^ in PBS; Merck Life Sciences, F0895), and maintained in growth medium supplemented with 10 ng/µL basic fibroblast growth factor (PeproTech, 100-18B). Plates were cultured in 5% CO2 at 37 °C and the media partially refreshed after 7 days. After 10 days, the cultures were washed in PBS, fixed in 10% neutral buffered formalin for 10 min at room temperature, washed again with sterile water, and stained with Harris hematoxylin for 5 min. Cells were imaged using an EVOS M7000 and analyzed in ImageJ using the cell counter plugin to enumerate clonal colonies comprising 50 cells or more.

### Enzyme-linked immunosorbent assay (ELISA)

Spent media of monolayer or assembloid cultures was collected and cleared of debris by centrifugation (16000 ×*g*, 10 min at 4°C). Quantitation of analytes in cell supernatant was performed using DuoSet sandwich ELISA kits exactly (Bio-Techne, Abingdon, UK). Standard curves were fitted to a 4-parameter logistic fit curve and analyte concentrations interpolated from these graphs.

---

**Fig. S1.**
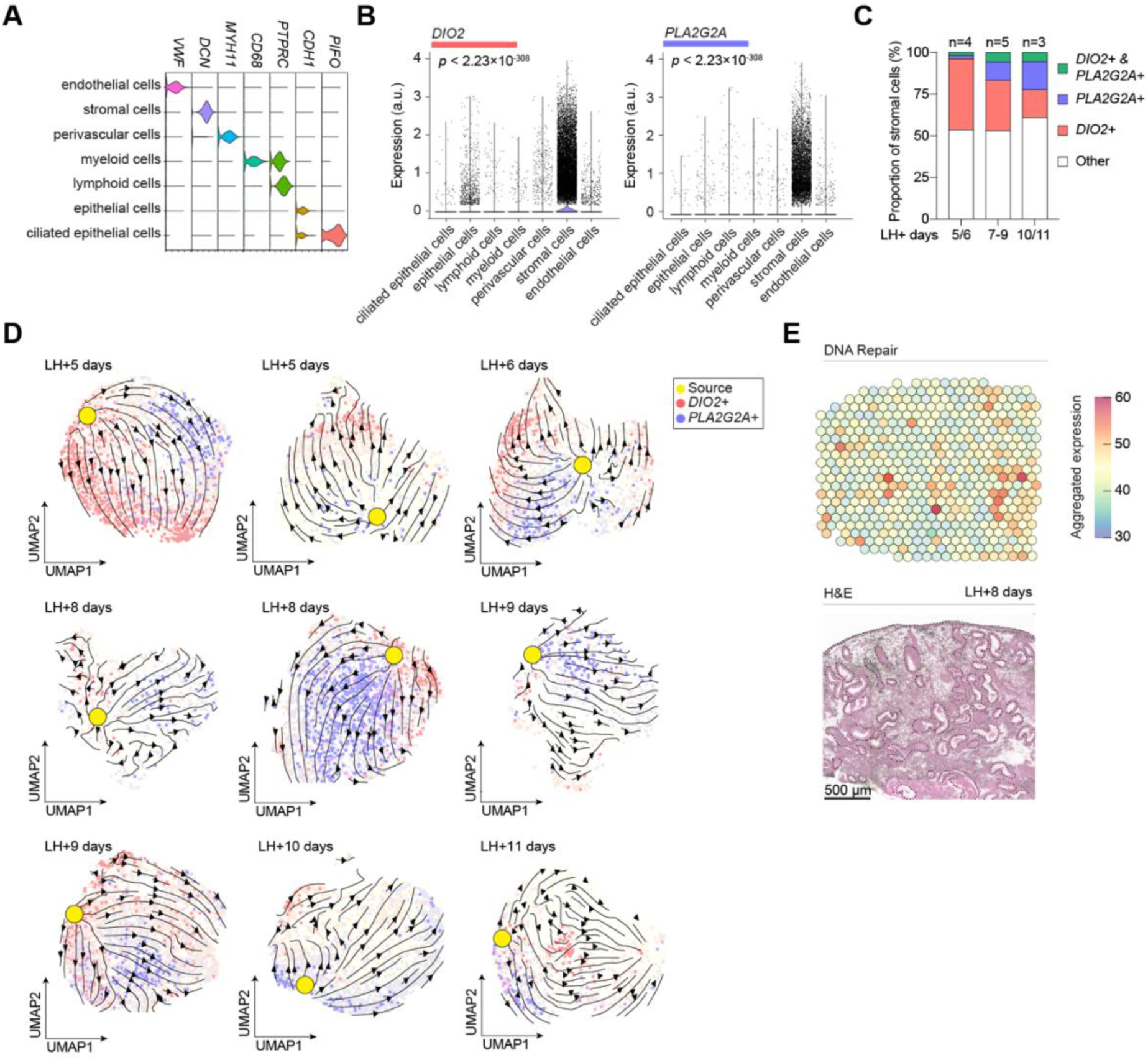
Single-cell transcriptomics of endometrial stromal subsets. (**A**) Violin plots showing expression of canonical marker genes in different endometrial cell types. (**B**) Violin plots showing *PLA2G2A* and *DIO2* expression across different endometrial cell types with *p*-values based on Wilcoxon rank sum test; a.u., arbitrary units. (**C**) Column bar graphs depicting the relative abundance of stromal subsets across the indicated cycle days. (**D**) RNA velocity mapped onto UMAP plots of stromal subsets in nine endometrial biopsies. (**E**) Visualization of the GSEA hallmark ‘DNA Repair’ gene set in mid-luteal endometrium by Visium spatial transcriptomics (upper panel). The lower panel shows the corresponding tissue section stained with H&E.

**Fig. S2.**
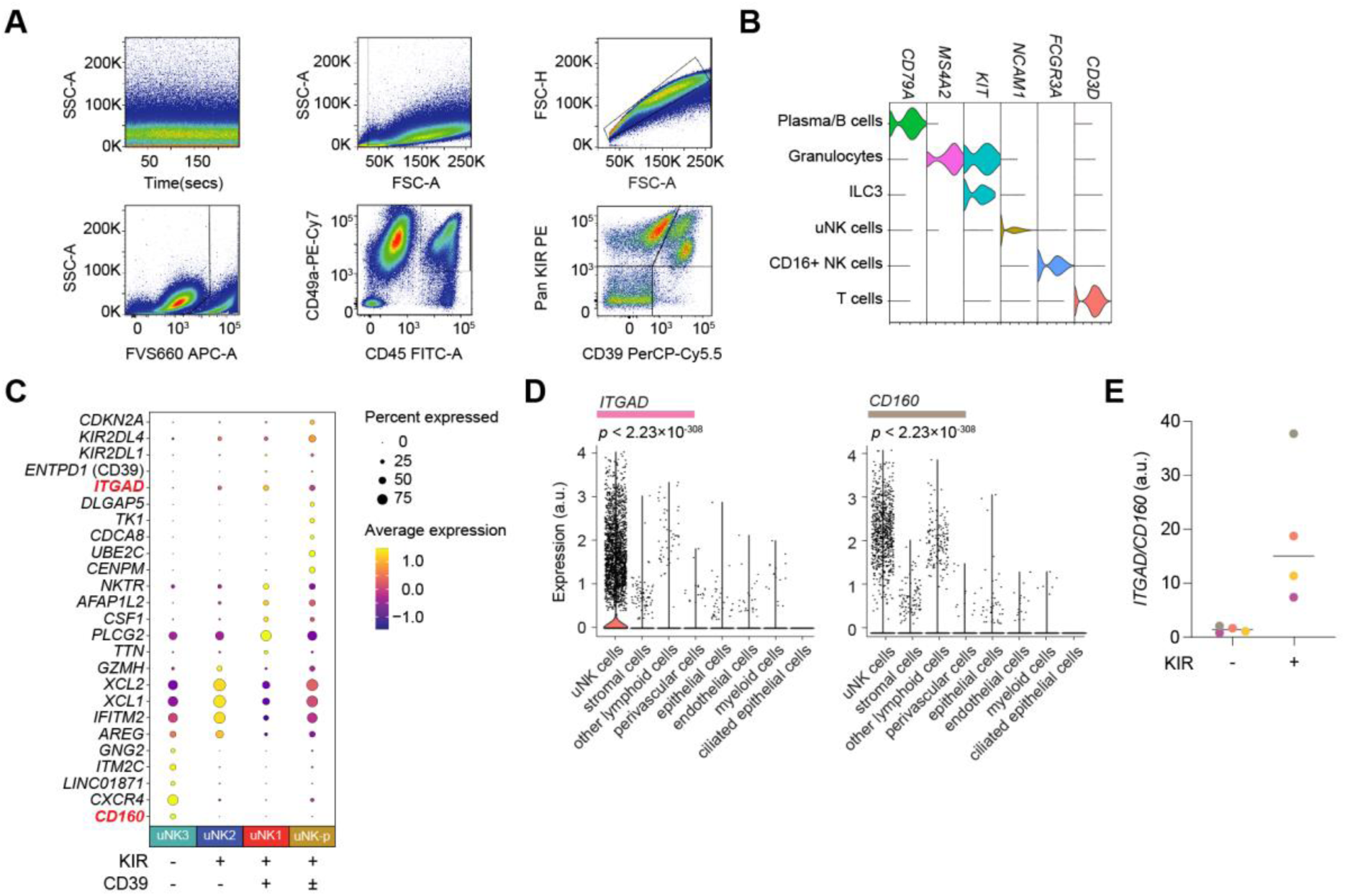
Analysis of uNK subsets. (**A**) Flow cytometry gating strategy for uNK cell subpopulations. (**B**) Violin plot based on scRNA-seq data of selected marker genes for different immune cell populations in peri-implantation endometrium. (**C**) Dot plot of expression levels of selected differentially expressed genes in uNK1-3 subsets; uNK-p denotes proliferating uNK cells. Dot size represents the percentage of positive cells, and the colour key indicates average gene expression levels. (**D**) Violin plots of *ITGAD* and *CD160* expression in different endometrial cell types. Statistical significance was determined using the Wilcoxon rank sum test. (**E**) Ratio of *ITGAD* and *CD160* transcripts in KIR+ and KIR-uNK subsets purified by FACS. *ITGAD* and *CD160* transcript levels were quantified by ddPCR. The data show individual ratios and median ratio from four biological repeat experiments; a.u., arbitrary units.

**Fig. S3.**
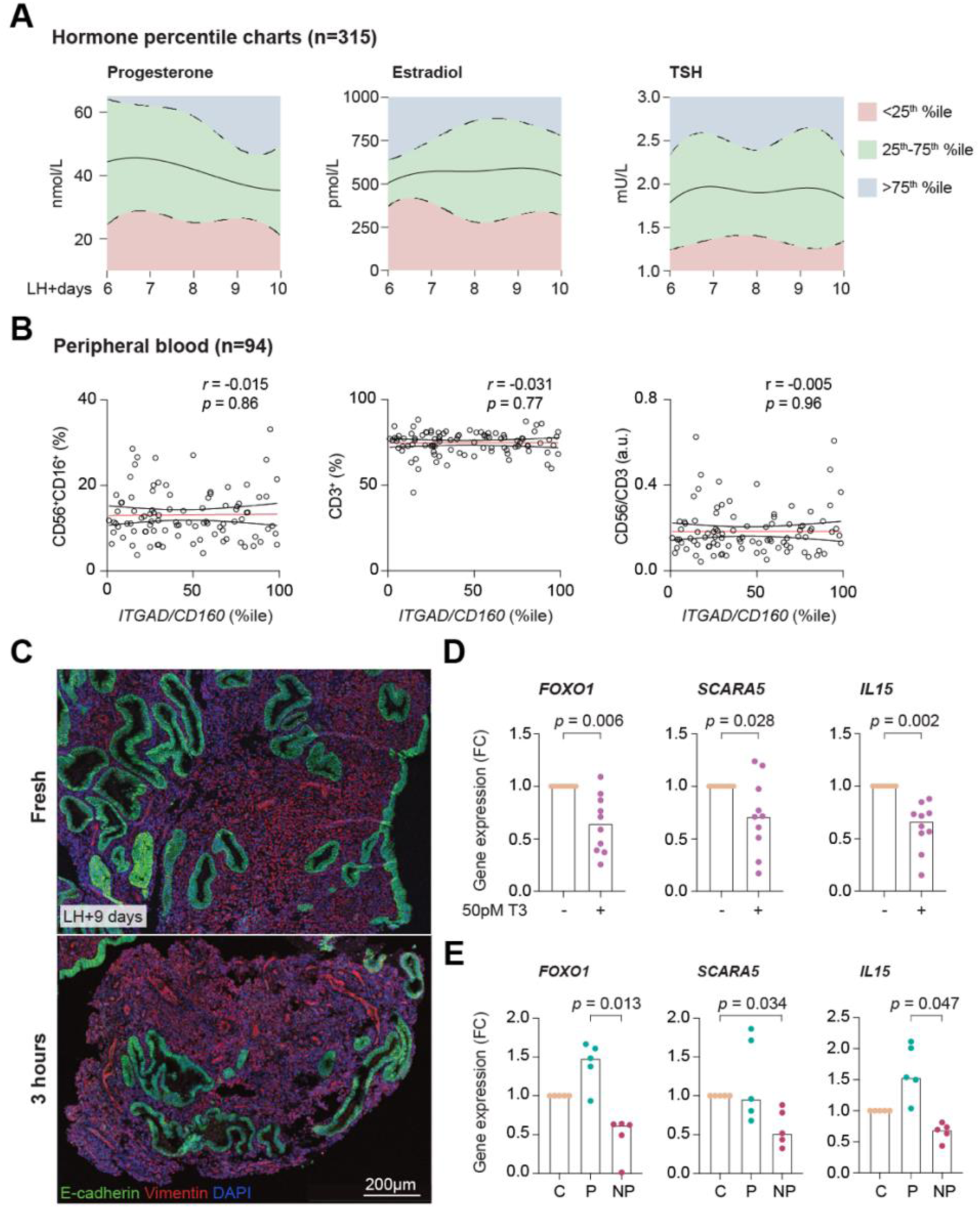
Hormones and peripheral blood immune cells. (**A**) Circulating levels of progesterone (nmol/L), estradiol (pmol/L) and TSH (mU/L) in 315 subjects 6 to 10 days after the LH surge. The solid line denotes median circulating levels, and the dotted line show upper and lower quartiles. (**B**) Spearman’s correlation between uNK cell expansion, measured by the percentile rank of normalized *ITGAD*/*CD160* ratios, and the relative abundance of circulating CD56+ NK cells, CD3+ T cells and CD56+/CD3+ cell ratios in paired peripheral blood and endometrial samples from 94 subjects. (**C**) Representative images of vimentin (red) and cadherin 1 (green) immunofluorescence in a freshly isolated tissue section before and after incubation in additive-free media (AFM). Nuclei were stained with DAPI. (**D**) Freshly isolated endometrial samples were divided, and sections incubated for 3 hours in AFM or media supplemented with 50 pM triiodothyronine (T3) prior to gene expression analysis by RT-qPCR. Bar graphs show fold change (FC) in the transcript levels of pre-decidual marker genes in response to T3 treatment in 10 biological repeat experiments. Individual data points are also shown. Statistical significance was determined using the Wilcoxon matched pairs signed rank test. (**E**) Freshly isolated endometrial samples were divided, and sections incubated for 3 hours in pooled spent medium of IVF blastocysts, resulting in pregnancy (P) or not (NP, not pregnant). Embryo-free droplets collected in parallel served as the control (c) group. Bar graphs showing median FC in the transcript levels of pre-decidual marker genes in five biological repeat experiments. Individual data points are also shown; *p*-values based on the Friedman and Dunn’s multiple comparison test.

**Fig. S4.**
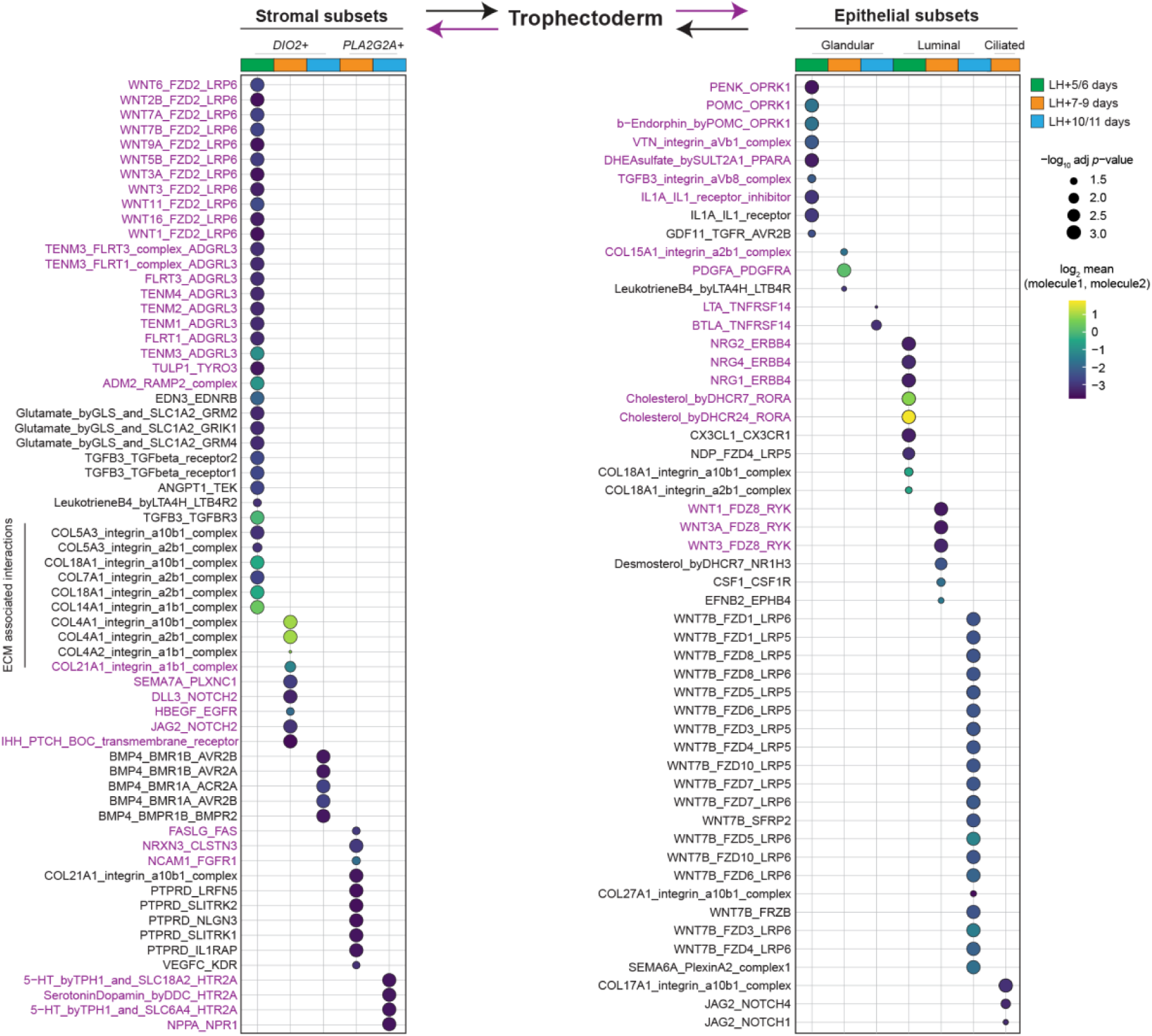
Embryonic trophectoderm-endometrium crosstalk. Dot plots showing CellPhoneDB predictions of receptor-ligand interactions of between endometrial stromal subsets (left panel) and epithelial cell types (right panel) cells and day-6/7 embryonic trophectoderm. Interaction involving embryonic trophectoderm and endometrial ligands are highlighted in purple and black lettering, respectively. Circle size and color scale indicate FDR-adjusted *p*-value and the mean of the average expression values of genes encoding interacting molecules, respectively.

**Fig. S5.**
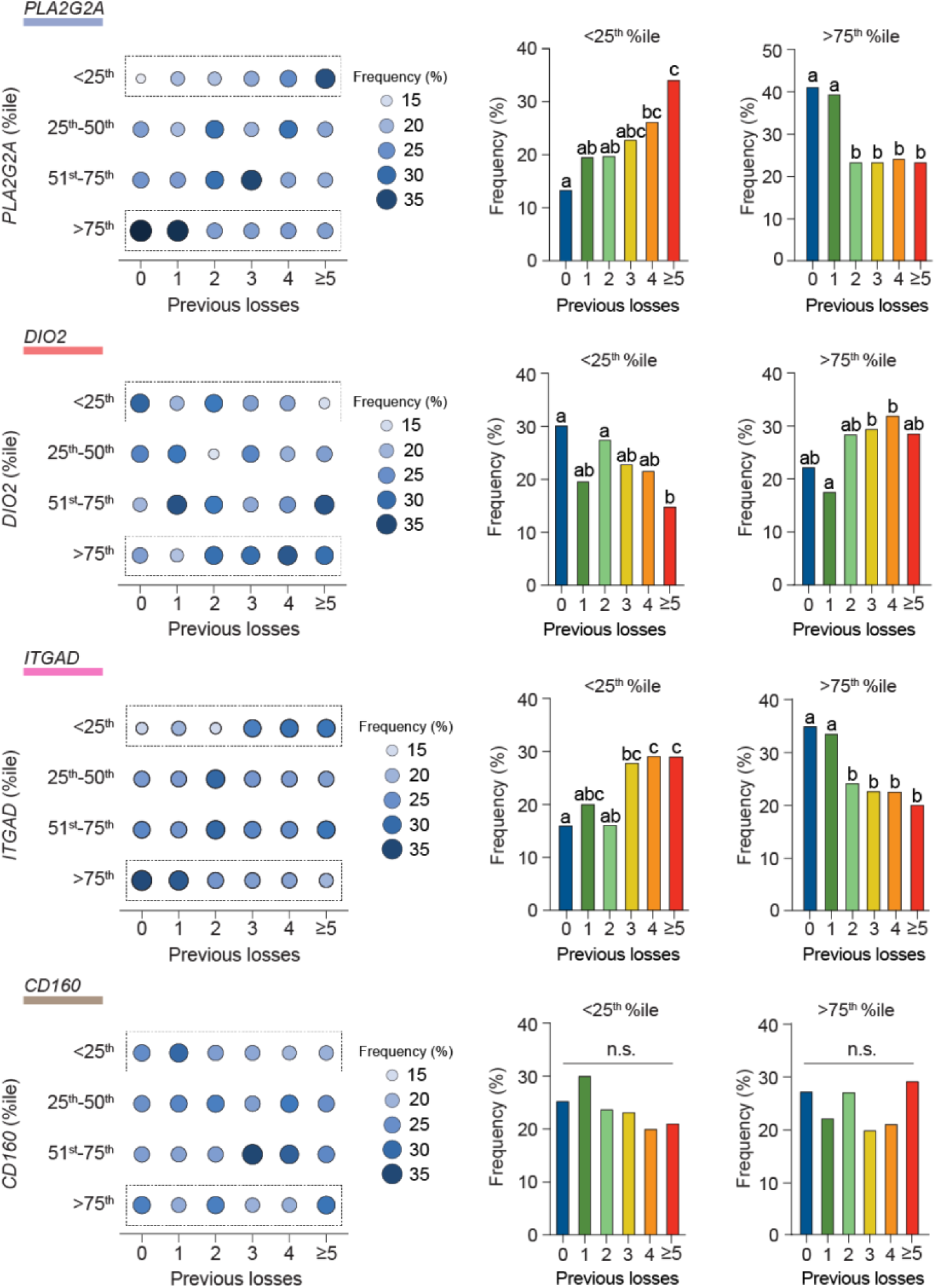
Stromal and uNK subset marker genes stratified by previous miscarriages. Dot plots of normalized marker genes grouped in quartile bins and stratified by the number of previous pregnancy losses (left panels). Dot sizes and color gradients denote the proportion (%) of subjects in each quartile. The number of subjects in each miscarriage group is also indicated. Bar charts (right panels) enumerating the proportion of subjects in the lowest (<25^th^ %ile) and highest (>75^th^ %ile) quartile bin. Different letters above the bars indicate groups that are significantly different from each other at *p* < 0.05, Chi-squared test with Bonferroni correction.

**Fig. S6.**
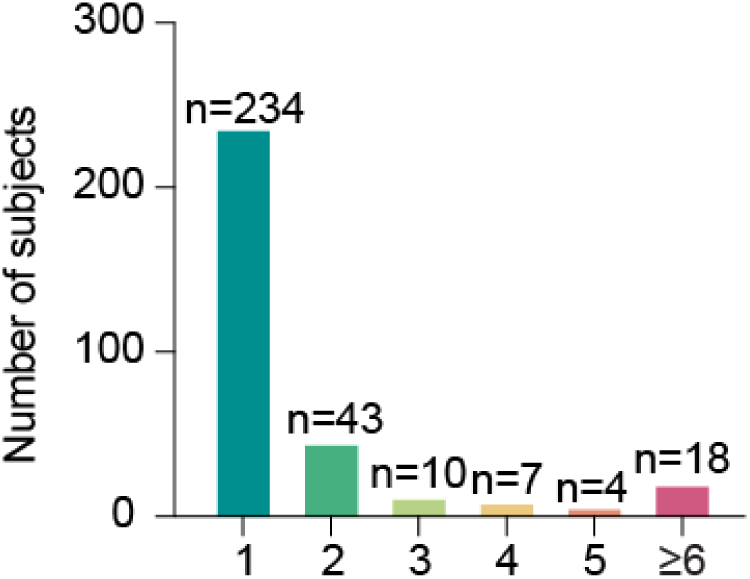
Number of cycles between A and B biopsies.

**Table S1.** Subject demographics for bulk RNA-seq analysis of paired endometrial biopsies.

| **Sample** | **Age (years)** | **LH+(days)** | **BMI** | **Days**<br>**between**<br>**biopsies** | ***SCARA5/DIO2* (%ile)** |
| --- | --- | --- | --- | --- | --- |
| X17476 | 42 | 10 | 20 | 28 | 4.5 |
| X17515 |  | 9 |  |  | 30.4 |
| X17524 | 37 | 9 | 29.5 | 28 | 82.4 |
| X17581 |  | 10 |  |  | 9.9 |
| X19257 | 29 | 9 | 23 | 56 | 39.6 |
| X19361 |  | 10 |  |  | 83.2 |
| X17728 | 37 | 8 | 22 | 102 | 57.1 |
| X18185 |  | 9 |  |  | 24.7 |
| X17153 | 32 | 10 | 28 | 24 | 44.5 |
| X17196 |  | 11 |  |  | 8.4 |
| X19233 | 38 | 7 | 23 | 115 | 25.9 |
| X19440 |  | 8 |  |  | 41.8 |

**Table S2.** Subject demographics for functional assays.

| **Assay** | **Figure** | **N=** | **Age (years)**<br>**median (IQR)** | **LH+(days)**<br>**median (IQR)** | **BMI**<br>**median (IQR)** |
| --- | --- | --- | --- | --- | --- |
| single-cell RNA-seq | 2F-H | 4 | 36 (32.5-40.3) | 5-6 | 25.8 (24-27.9) |
|  |  | 5 | 35 (32-39) | 7-9 | 24.0 (23-25) |
|  |  | 3 | 38 (30-42) | 10-11 | 29 (28-29) |
| uNK cell flow<br>cytometry | 2B-2C | 6 | 39 (34-42) | 6 | 25 (23-29.5) |
|  |  | 10 | 37.5 (32-40.25) | 7 | 23 (22-27) |
|  |  | 12 | 36.5 (33-39.75) | 8 | 23 (22-25.8) |
|  |  | 11 | 35 (32-40) | 9 | 24 (23.25-29.8) |
|  |  | 12 | 34.5 (33-39.75) | 10 | 25 (23.25-27) |
|  |  | 4 | 39.5 (32-45.5) | 11 | 23 (21.25-27.8) |
| FACS (uNK subsets -<br>killing assay) | 2E | 8 | 35 (34-38) | 7 (5-7.8) | 26 (21.8-29.8) |
| primary stromal cell<br>cultures |  | 10 | 35.5 (33.5-36) | 8.5 (8-9) | 24 (23-27) |
| FACS (uNK subsets -<br>purity) |  | 11 | 38 (32.75-40) | 8 (7-9) | 26.5 (23.3-30) |
| Quantification of<br>marker genes by<br>ddPCR | 2I-J | 57 | 37 (34-38) | 6 | 24 (21-26.5) |
|  |  | 179 | 35 (33-37) | 7 | 24 (22-29) |
|  |  | 207 | 35 (32-38) | 8 | 24 (21-27) |
|  |  | 202 | 34 (32-37) | 9 | 24 (22-27) |
|  |  | 134 | 36 (33-38.5) | 10 | 25 (22.6-28) |
| Spatial analysis of<br>marker genes | 3C, E, F<br>&<br>fig.<br>S3A-C | 264 | 36 (33-39) | 8 (7-9) | 24 (22-28) |
| Circulating hormone<br>levels |  | 315 | 36 (33-39) | 8 (7-9) | 24 (22-27.5) |
| Peripheral blood<br>immune cells |  | 94 | 36 (33-39) | 8 (7-9) | 24.24 (21.8-27) |
| T3 experiment | 3H, K | 11 | 35 (33-39) | 8 (8-9) | 22 (21-26) |
| Blastocyst conditioned<br>media experiment |  | 5 | 36 (34-39.5) | 8 (7-8.25) | 20 (20.7-31) |
| Prolonged<br>decidualization | 6C-G | 5 | 33 (28-38.5) | 8 (7-9.5) | 24 (21-31) |
| Cyclical<br>decidualization | 6H-I | 7 | 35.5 (35-37) | 8.5 (6.5-10.25) | 32 (27.5-35.5) |

**Table S3.**
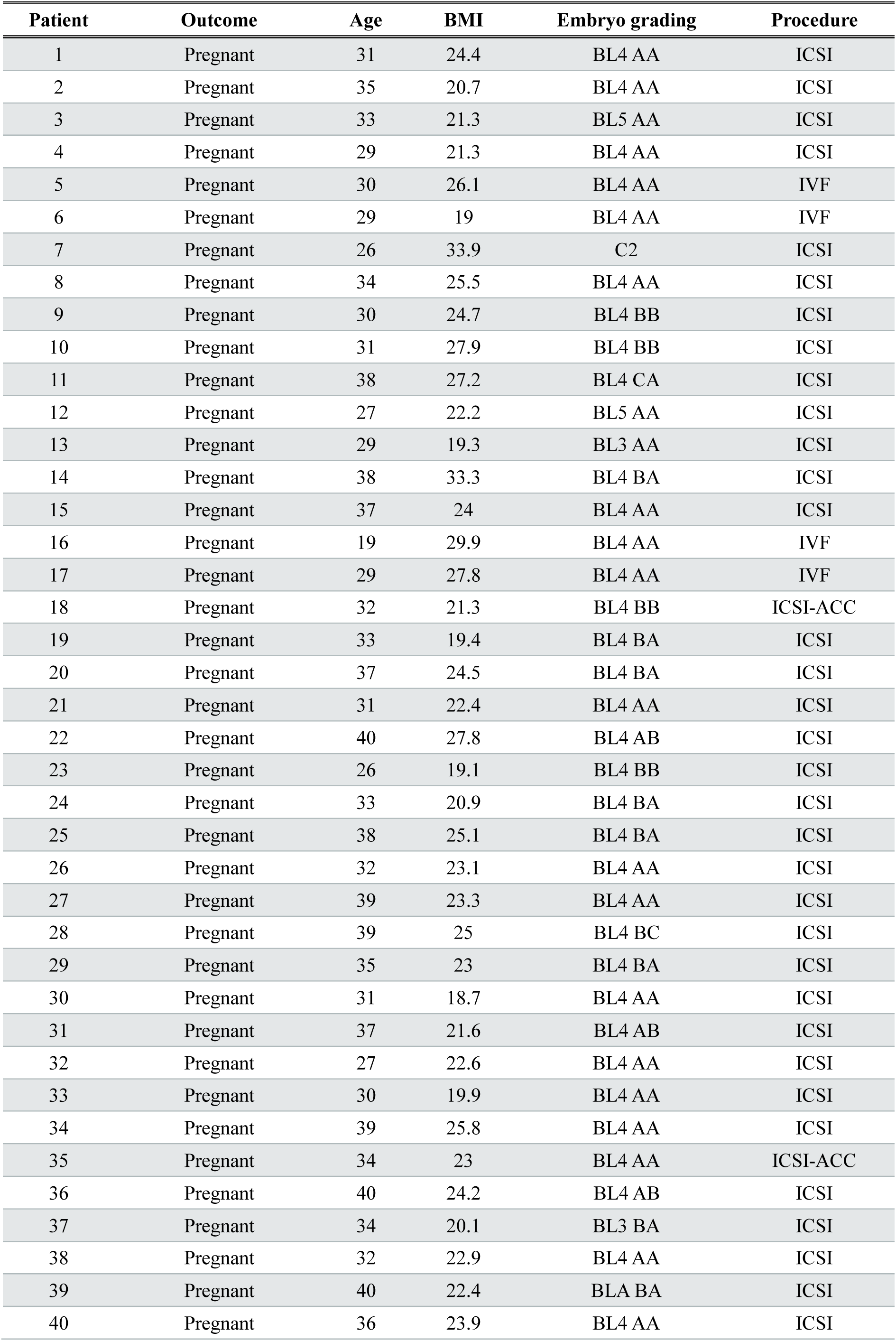
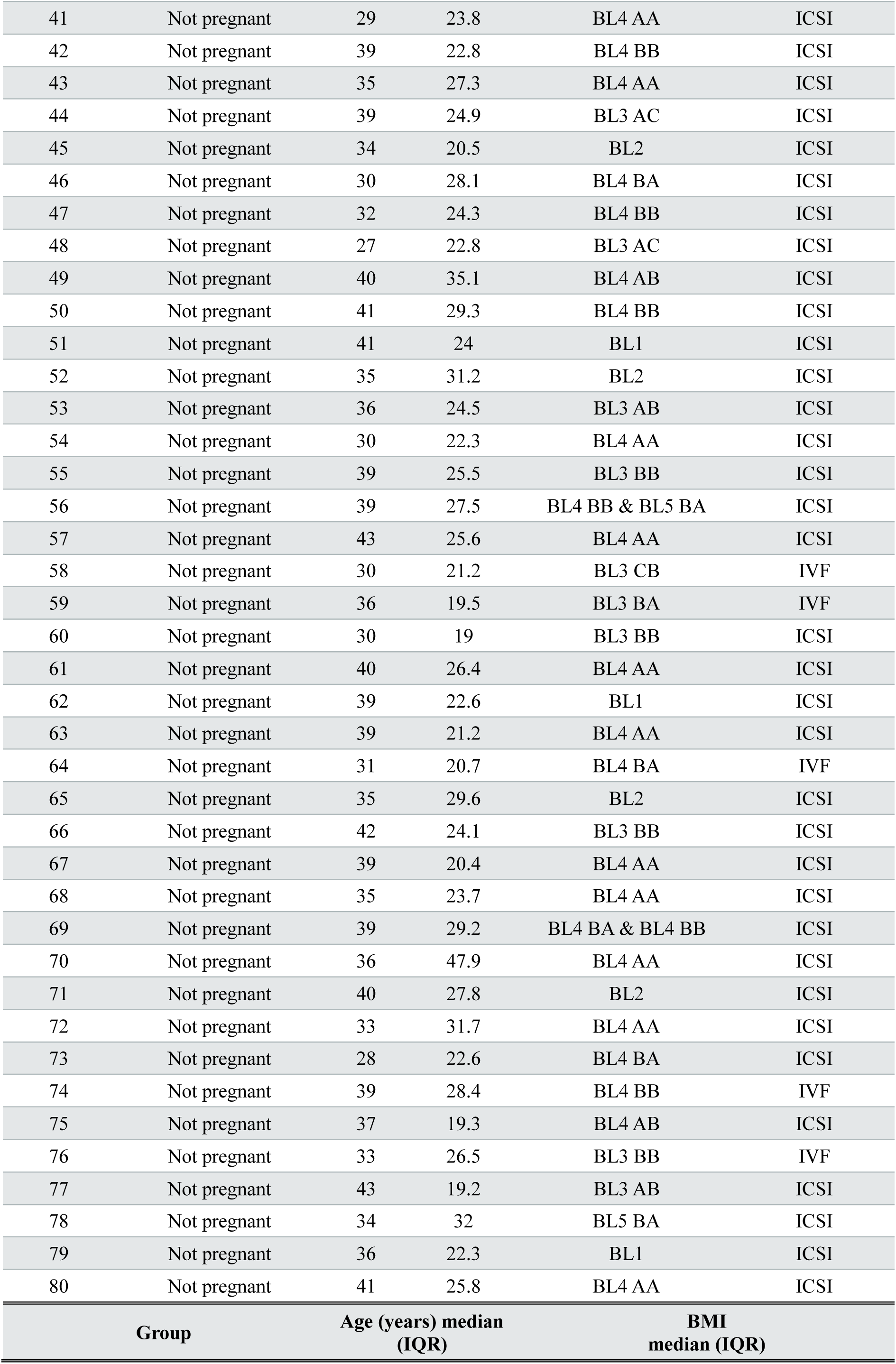
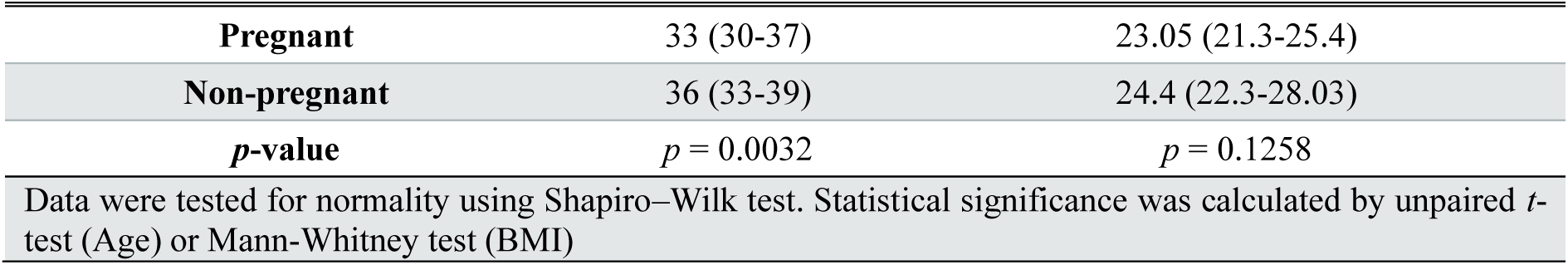
Embryo grading and subject demographics.

| Patient | Outcome | Age | BMI | Embryo grading | Procedure |
| --- | --- | --- | --- | --- | --- |
| 1 | Pregnant | 31 | 24.4 | BL4 AA | ICSI |
| 2 | Pregnant | 35 | 20.7 | BL4 AA | ICSI |
| 3 | Pregnant | 33 | 21.3 | BL5 AA | ICSI |
| 4 | Pregnant | 29 | 21.3 | BL4 AA | ICSI |
| 5 | Pregnant | 30 | 26.1 | BL4 AA | IVF |
| 6 | Pregnant | 29 | 19 | BL4 AA | IVF |
| 7 | Pregnant | 26 | 33.9 | C2 | ICSI |
| 8 | Pregnant | 34 | 25.5 | BL4 AA | ICSI |
| 9 | Pregnant | 30 | 24.7 | BL4 BB | ICSI |
| 10 | Pregnant | 31 | 27.9 | BL4 BB | ICSI |
| 11 | Pregnant | 38 | 27.2 | BL4 CA | ICSI |
| 12 | Pregnant | 27 | 22.2 | BL5 AA | ICSI |
| 13 | Pregnant | 29 | 19.3 | BL3 AA | ICSI |
| 14 | Pregnant | 38 | 33.3 | BL4 BA | ICSI |
| 15 | Pregnant | 37 | 24 | BL4 AA | ICSI |
| 16 | Pregnant | 19 | 29.9 | BL4 AA | IVF |
| 17 | Pregnant | 29 | 27.8 | BL4 AA | IVF |
| 18 | Pregnant | 32 | 21.3 | BL4 BB | ICSI-ACC |
| 19 | Pregnant | 33 | 19.4 | BL4 BA | ICSI |
| 20 | Pregnant | 37 | 24.5 | BL4 BA | ICSI |
| 21 | Pregnant | 31 | 22.4 | BL4 AA | ICSI |
| 22 | Pregnant | 40 | 27.8 | BL4 AB | ICSI |
| 23 | Pregnant | 26 | 19.1 | BL4 BB | ICSI |
| 24 | Pregnant | 33 | 20.9 | BL4 BA | ICSI |
| 25 | Pregnant | 38 | 25.1 | BL4 BA | ICSI |
| 26 | Pregnant | 32 | 23.1 | BL4 AA | ICSI |
| 27 | Pregnant | 39 | 23.3 | BL4 AA | ICSI |
| 28 | Pregnant | 39 | 25 | BL4 BC | ICSI |
| 29 | Pregnant | 35 | 23 | BL4 BA | ICSI |
| 30 | Pregnant | 31 | 18.7 | BL4 AA | ICSI |
| 31 | Pregnant | 37 | 21.6 | BL4 AB | ICSI |
| 32 | Pregnant | 27 | 22.6 | BL4 AA | ICSI |
| 33 | Pregnant | 30 | 19.9 | BL4 AA | ICSI |
| 34 | Pregnant | 39 | 25.8 | BL4 AA | ICSI |
| 35 | Pregnant | 34 | 23 | BL4 AA | ICSI-ACC |
| 36 | Pregnant | 40 | 24.2 | BL4 AB | ICSI |
| 37 | Pregnant | 34 | 20.1 | BL3 BA | ICSI |
| 38 | Pregnant | 32 | 22.9 | BL4 AA | ICSI |
| 39 | Pregnant | 40 | 22.4 | BLA BA | ICSI |
| 40 | Pregnant | 36 | 23.9 | BL4 AA | ICSI |

| 41 | Not pregnant | 29 | 23.8 | BL4 AA | ICSI |
| --- | --- | --- | --- | --- | --- |
| 42 | Not pregnant | 39 | 22.8 | BL4 BB | ICSI |
| 43 | Not pregnant | 35 | 27.3 | BL4 AA | ICSI |
| 44 | Not pregnant | 39 | 24.9 | BL3 AC | ICSI |
| 45 | Not pregnant | 34 | 20.5 | BL2 | ICSI |
| 46 | Not pregnant | 30 | 28.1 | BL4 BA | ICSI |
| 47 | Not pregnant | 32 | 24.3 | BL4 BB | ICSI |
| 48 | Not pregnant | 27 | 22.8 | BL3 AC | ICSI |
| 49 | Not pregnant | 40 | 35.1 | BL4 AB | ICSI |
| 50 | Not pregnant | 41 | 29.3 | BL4 BB | ICSI |
| 51 | Not pregnant | 41 | 24 | BL1 | ICSI |
| 52 | Not pregnant | 35 | 31.2 | BL2 | ICSI |
| 53 | Not pregnant | 36 | 24.5 | BL3 AB | ICSI |
| 54 | Not pregnant | 30 | 22.3 | BL4 AA | ICSI |
| 55 | Not pregnant | 39 | 25.5 | BL3 BB | ICSI |
| 56 | Not pregnant | 39 | 27.5 | BL4 BB & BL5 BA | ICSI |
| 57 | Not pregnant | 43 | 25.6 | BL4 AA | ICSI |
| 58 | Not pregnant | 30 | 21.2 | BL3 CB | IVF |
| 59 | Not pregnant | 36 | 19.5 | BL3 BA | IVF |
| 60 | Not pregnant | 30 | 19 | BL3 BB | ICSI |
| 61 | Not pregnant | 40 | 26.4 | BL4 AA | ICSI |
| 62 | Not pregnant | 39 | 22.6 | BL1 | ICSI |
| 63 | Not pregnant | 39 | 21.2 | BL4 AA | ICSI |
| 64 | Not pregnant | 31 | 20.7 | BL4 BA | IVF |
| 65 | Not pregnant | 35 | 29.6 | BL2 | ICSI |
| 66 | Not pregnant | 42 | 24.1 | BL3 BB | ICSI |
| 67 | Not pregnant | 39 | 20.4 | BL4 AA | ICSI |
| 68 | Not pregnant | 35 | 23.7 | BL4 AA | ICSI |
| 69 | Not pregnant | 39 | 29.2 | BL4 BA & BL4 BB | ICSI |
| 70 | Not pregnant | 36 | 47.9 | BL4 AA | ICSI |
| 71 | Not pregnant | 40 | 27.8 | BL2 | ICSI |
| 72 | Not pregnant | 33 | 31.7 | BL4 AA | ICSI |
| 73 | Not pregnant | 28 | 22.6 | BL4 BA | ICSI |
| 74 | Not pregnant | 39 | 28.4 | BL4 BB | IVF |
| 75 | Not pregnant | 37 | 19.3 | BL4 AB | ICSI |
| 76 | Not pregnant | 33 | 26.5 | BL3 BB | IVF |
| 77 | Not pregnant | 43 | 19.2 | BL3 AB | ICSI |
| 78 | Not pregnant | 34 | 32 | BL5 BA | ICSI |
| 79 | Not pregnant | 36 | 22.3 | BL1 | ICSI |
| 80 | Not pregnant | 41 | 25.8 | BL4 AA | ICSI |
| Group | Age (years) median (IQR) |  | BMI median (IQR) |  |  |

|  |  |  |
| --- | --- | --- |
| **Pregnant** | 33 (30-37) | 23.05 (21.3-25.4) |
| **Non-pregnant** | 36 (33-39) | 24.4 (22.3-28.03) |
| ***p*-value** | *p* = 0.0032 | *p* = 0.1258 |
| Data were tested for normality using Shapiro–Wilk test. Statistical significance was calculated by unpaired *t*-test (Age) or Mann-Whitney test (BMI) |  |  |

**Table S4.** Subject demographics for retrospective cohort.

| **Number of miscarriages**<br>**median (range)** | **N=** | **Age (years)**<br>**median (IQR)** | **LH+ (days)**<br>**median (IQR)** | **BMI**<br>**median (IQR)** | **Live births**<br>**mean [range]** |
| --- | --- | --- | --- | --- | --- |
| 0 | 162 | 35<br>(33-38) | 8<br>(7-9) | 23<br>(21-26) | 0.09<br>[0-2] |
| 1 | 99 | 36<br>(33-38) | 8<br>(7-10) | 24<br>(22-27) | 0.11<br>[0-2] |
| 2 | 92 | 36<br>(32-38.75) | 8<br>(7-10) | 24<br>(21-26.8) | 0.12<br>[0-1] |
| 3 | 88 | 35<br>(32-38) | 8<br>(8-10) | 24<br>(22-28)* | 0.18<br>[0-3] |
| 4 | 86 | 34.5<br>(32.8-36.3) | 8<br>(7-9) | 24<br>(22-27) | 0.20<br>[0-2] |
| ≥5<br>(5-18) | 136 | 35<br>(32-38) | 8<br>(7-10) | 25<br>(23-29)* | 0.33<br>[0-3]* |
| Data were tested for normality using Shapiro–Wilk test. Statistical significance was calculated by ANOVA with Dunnett's multiple comparison test for normally distributed data (age) or Kruskal-Wallis test with Dunn's multiple comparison test for non-normally distributed data (BMI, LH+, Previous miscarriages and live births) *denotes $p < 0.05$ compared to subjects with <u>no</u> previous pregnancy loss. | | | | | |

**Table S5.**
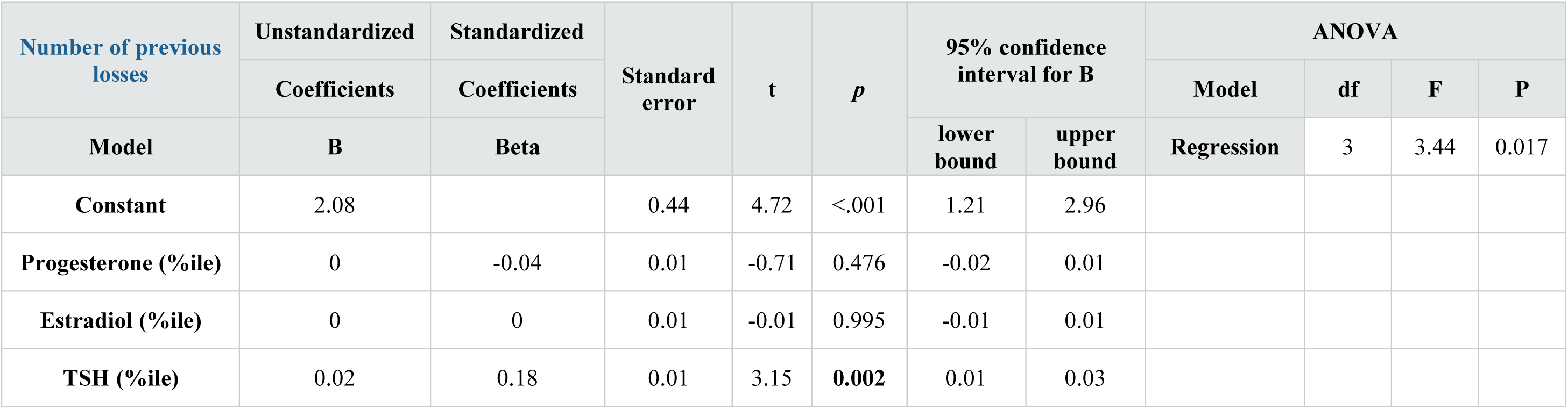
Multilinear regression analysis: Number of miscarriages and normalized circulating hormone levels.

**Table S6.** Multilinear regression analysis: marker gene ratios.

| *PLA2G2A/DIO2*<br>(%ile) | Unstandardize<br>d | Standardized | Standard<br>error | t | p | 95% confidence<br>interval for B |  | ANOVA |  |  |  |
| --- | --- | --- | --- | --- | --- | --- | --- | --- | --- | --- | --- |
|  | Coefficients | Coefficients |  |  |  |  |  | Model | df | F | p |
|  | B | Beta |  |  |  | lower<br>bound | upper<br>bound | Regression | 6 | 15.26 | <.001 |
| Constant | 57.57 |  | 12.71 | 4.53 | <.001 | 32.51 | 82.63 |  |  |  |  |
| Number of previous<br>losses | -1.36 | -0.13 | 0.42 | -3.23 | **0.001** | -2.19 | -0.53 |  |  |  |  |
| Age | -0.3 | -0.04 | -1.11 | -1.11 | 0.269 | -0.82 | 0.23 |  |  |  |  |
| Live births | 1.6 | 0.02 | 0.62 | 0.62 | 0.538 | -3.52 | 6.72 |  |  |  |  |
| BMI | -0.48 | -0.08 | 0.24 | -1.97 | 0.05 | -0.96 | 0 |  |  |  |  |
| *ITGAD/CD160*<br>(%ile) | Unstandardize<br>d | Standardized | Standard<br>error | t | p | 95% confidence<br>interval for B |  | ANOVA |  |  |  |
|  | Coefficients | Coefficients |  |  |  |  |  | Model | df | F | p |
|  | B | Beta |  |  |  | lower<br>bound | upper<br>bound | Regression | 6 | 14.34 | <.001 |
| Constant | 52.73 |  | 12.82 | 4.11 | <.001 | 27.45 | 78 |  |  |  |  |
| Number of previous<br>losses | -1.23 | -0.12 | 0.42 | -2.91 | **0.004** | -2.07 | -0.4 |  |  |  |  |
| Age | -0.06 | -0.01 | 0.27 | -0.24 | 0.811 | -0.6 | 0.47 |  |  |  |  |
| Live births | -1.94 | -0.03 | 2.61 | -0.74 | 0.458 | -7.09 | 3.21 |  |  |  |  |
| BMI | -0.29 | -0.05 | 0.25 | -1.18 | 0.238 | -0.78 | 0.2 |  |  |  |  |

**Table S7.**
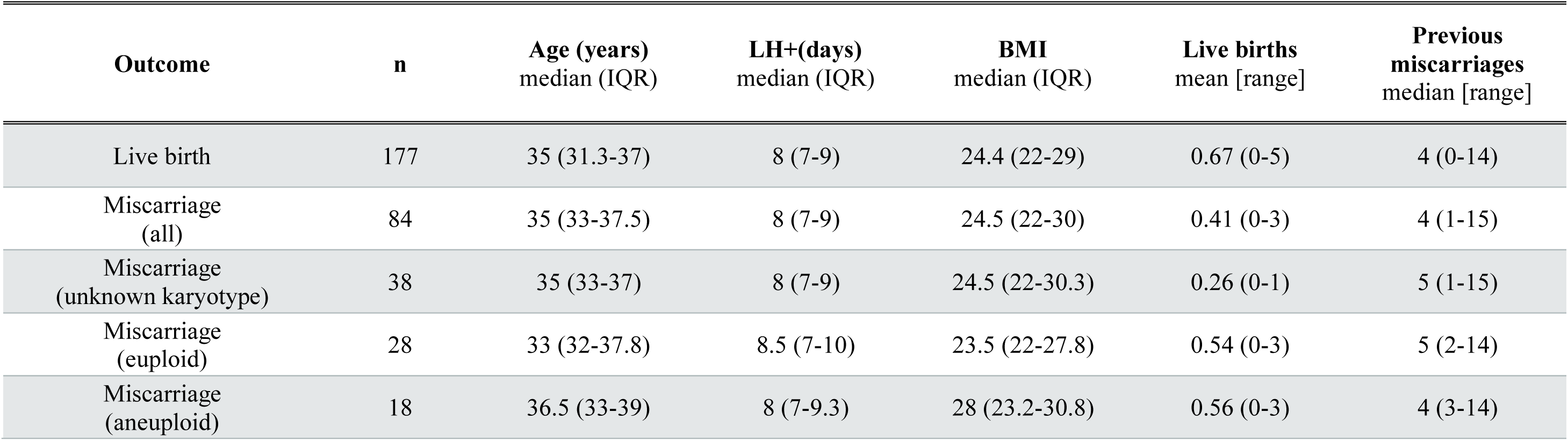
Subject demographics for prospective cohort.

**Table S8.**
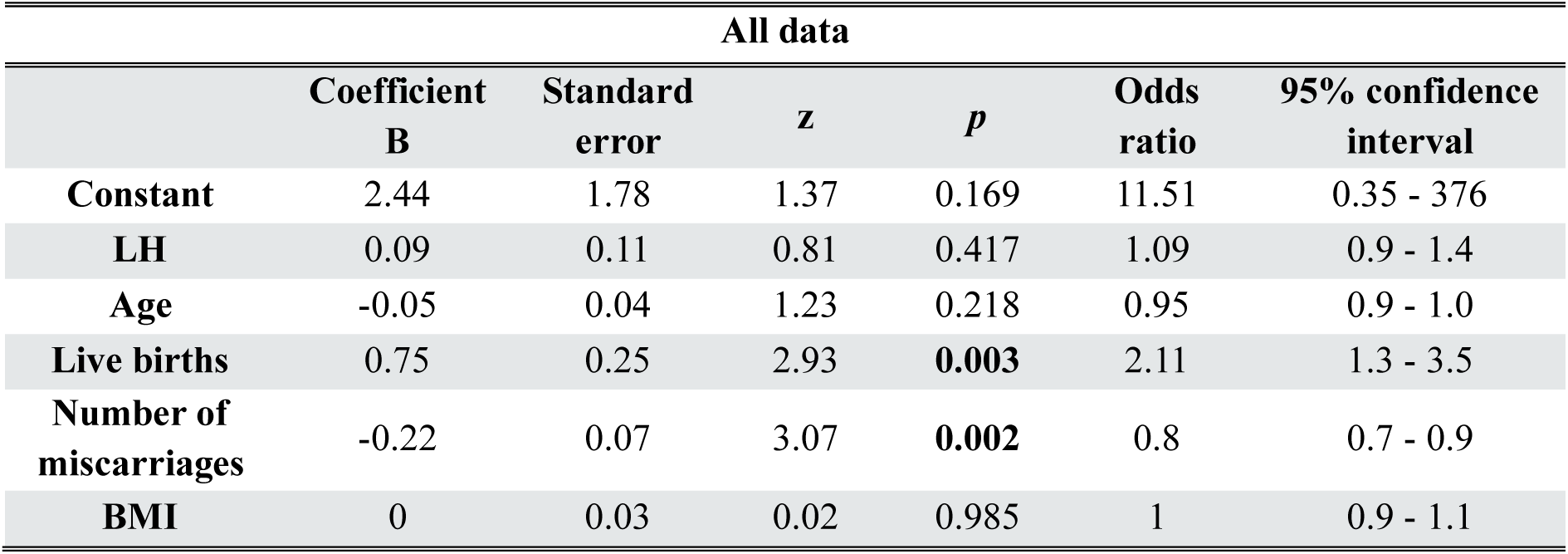
Logistic regression analysis for prospective cohort.

**Table S9.** Subject demographics for paired endometrial biopsy analysis.

| **Group**<br>(number of<br>previous<br>miscarriage) | **N=** | **Previous**<br>**miscarriages**<br>median (range) | **Age**<br>**(years)**<br>median<br>(IQR) | **LH+(days)**<br>median<br>(IQR) | **BMI**<br>median (IQR) | **Live**<br>**births**<br>mean<br>[range] | **Days**<br>**between**<br>**biopsies**<br>median<br>(IQR) |
| --- | --- | --- | --- | --- | --- | --- | --- |
| 0 | 93 | 0 | 35 (32-39) | 8 (7-9) | 23 (21-26) | 0.1 [0-2] | 32 (28-56) |
| 1-2 | 90 | 2 (1-2) | 35 (32-39) | 8 (7-9) | 24.5 (21-28) | 0.1 [0-2] | 31 (28-57) |
| 3-4 | 85 | 3 (3-4) | 34 (32-36) | 9 (7-9) | 24 (22-28) | 0.3 [0-2]* | 31 (28-53) |
| ≥5 | 48 | 5 (5-18) | 35 (33-38) | 8.5 (7-10) | 27 (23-30)* | 0.5 [0-4]* | 28 (28-35) |
| Data were tested for normality using Shapiro–Wilk test. Statistical significance was calculated by ANOVA with Dunnett's multiple comparison test for normally distributed data (age) or Kruskal-Wallis test with Dunn's multiple comparison test for non-normally distributed data (BMI, LH+, previous miscarriages and live births). *denotes $p < 0.05$ compared to subjects with no previous pregnancy loss. | | | | | | | |

**Table S10.** Composition of assembloid culture media.

| **Expansion Medium (ExM)** |  |  |  |
| --- | --- | --- | --- |
| **Component** | **Manufacturer** | **Catalogue number** | **Final Concentration** |
| **Advanced DMEM/F12** | Life Technologies | 12634010 | 1× |
| **N2** | Life Technologies | 17502048 | 1× |
| **B27** | Life Technologies | 12587010 | 1× |
| **Antibiotic-Antimycotic** | Gibco | 15240062 | 1× |
| **L-glutamine** | Gibco | 25030081 | 2 mM |
| **A83-01** | Merck Life Science | SML0788-5MG | 500 nM |
| **R-spondin-1** | Peprtech | 120-38 | 500 ng/mL |
| **Nicotinamide** | Merck Life Science | N0636 | 10 mM |
| **N-acetyl-L-cysteine** | Merck Life Science | A9165-5G | 1.25 mM |
| **Noggin** | Peprtech | 120-10c | 100 ng/mL |
| **FGF10** | Peprtech | 100-26 | 100 ng/mL |
| **HGF** | Peprtech | 100-39 | 50 ng/mL |
| **EGF** | Peprtech | AF-100-15 | 50 ng/mL |
| **Minimal Decidualization Medium (MDM)** |  |  |  |
| **Component** | **Manufacturer** | **Catalogue number** | **Final Concentration** |
| **Advanced DMEM/F12** | Life Technologies | 12634010 | 1× |
| **N2** | Life Technologies | 17502048 | 1× |
| **B27** | Life Technologies | 12587010 | 1× |
| **Antibiotic-Antimycotic** | Gibco | 15240062 | 1× |
| **L-glutamine** | Gibco | 25030081 | 2 mM |
| **N-acetyl-L-cysteine** | Merck Life Science | A9165-5G | 1.25 mM |
| **8-bromo-cAMP** | Merck Life Science | B7880 | 0.5 mM |
| **MPA** | Merck Life Science | M1629 | 1 μM |
| **β-estradiol** | Merck Life Science | E2758 | 1 nM |

**Table S11.**
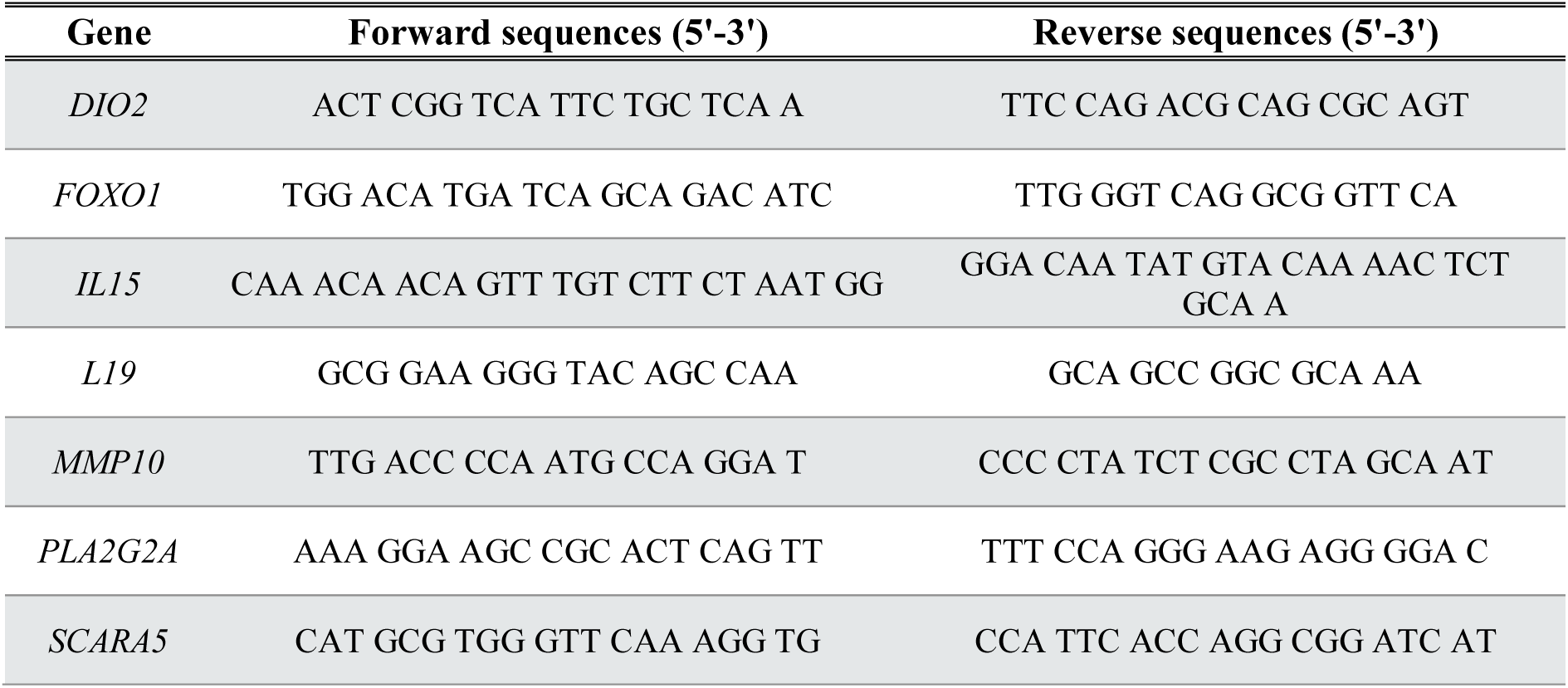
Primer sequences for RT-qPCR.

**Table S12.** Probes for ddPCR.

| Assay name | Assay type | Bio-Rad assay ID | Probe fluorophore |
| --- | --- | --- | --- |
| CD160 | Gene expression | dHsaCP2506689 | HEX |
| *DIO2* | Gene expression | dHsaCNS365622194 | HEX |
| *ITGAD* | Gene expression | dHsaCP2506311 | FAM |
| *PLA2G2A* | Gene expression | dHsaCPE5042784 | FAM |

**Table S13.** Probes for smISH.

| Gene | Manufacturer | Catalogue number | Channel |
| --- | --- | --- | --- |
| *DIO2* | Bio-Techne LTD | 562211 | C2 |
| *PLA2G2A* | Bio-Techne LTD | 581101 | C1 |

**Table S14.** Antibodies for immunohistochemistry.

| Antibody | Company | Catalogue number | Sample | Host species | Antigen retrieval buffer | Dilution |
| --- | --- | --- | --- | --- | --- | --- |
| CD56 | Leica Biosystems | NCL-LCD56-504 | Endometrial biopsy | Mouse | Tris-EDTA buffer, pH 9.0 | 1:100 in TBST |
| KIR2DL1/L2/L3/S1/S2 | Abcam | ab252938 | Endometrial biopsy | Rabbit | Tris-EDTA buffer, pH 9.0 | 1:1000 in TBST |
| E-cadherin | Cell Signalling Technology | 24E10 | Endometrial biopsy/explant culture | Rabbit | Sodium citrate buffer, pH 6.0 | 1:200 0.05% TBS-T |
| Vimentin | Cell Signalling Technology | 5G3F10 | Endometrial biopsy/explant culture | Mouse | Sodium citrate buffer, pH 6.0 | 1:200 0.05% TBST |
| E-cadherin | Cell Signalling Technology | 24E10 | IHC for assembloids | Rabbit | NA | 1:200 10%FBS-2%BSA-PBS-0.05%Tween-20 |
| Vimentin | Cell Signalling Technology | 5G3F10 | IHC for assembloids | Mouse | NA | 1:200 10%FBS-2%BSA-PBS-0.05%Tween-20 |
| Goat anti-Rabbit IgG Highly Cross-Absorbed Secondary Antibody, Alexa Fluor™ 647 | Invitrogen | A21245 | IHC for assembloids | Goat | NA | 1:1000 in 10%FBS-2%BSA-PBS-0.05%Tween-20 |
| Donkey anti-Mouse IgG Highly Cross-Absorbed Secondary Antibody, Alexa Fluor™ 594 | Invitrogen | A21203 | IHC for assembloids | Donkey | NA | 1:1000 in 10%FBS-2%BSA-PBS-0.05%Tween-20 |
| Donkey anti-Rabbit IgG (H+L) Highly Cross-Absorbed Secondary Antibody, Alexa Fluor™ 488 | Invitrogen | A21206 | Endometrial biopsy/explant culture | Donkey | NA | 1:1000 in 0.05% TBS-T |
| Donkey anti-Mouse IgG Highly Cross-Absorbed Secondary Antibody, Alexa Fluor™ 594 | Invitrogen | A21203 | Endometrial biopsy/explant culture | Donkey | NA | 1:1000 in 0.05% TBS-T |

**Table S15.** Antibodies for flow cytometry and sorting of primary uNK cells.

| **Antibody** | **Manufacturer** | **Catalogue number** | **Sample** | **Fluorophore** | **Clone** | **Dilution** |
| --- | --- | --- | --- | --- | --- | --- |
| Fixable Viability Stain 660 | BD Biosciences | 564405 | Endometrial tissue | APC | NA | 1:1000 in PBS |
| CD45 | BioLegend | 304006 | Endometrial tissue | FITC | HI30 | 1:40 in 0.5%BSA/PBS |
| CD49a | BioLegend | 328312 | Endometrial tissue | PE-Cy7 | TS2/7 | 1:40 in 0.5%BSA/PBS |
| CD158d (KI2DL4) | BioLegend | 347006 | Endometrial tissue | PE | MaB 33 | 1:40 in 0.5%BSA/PBS |
| CD158b (KIR2DL1/S1/S3/S5) | BioLegend | 339506 | Endometrial tissue | PE | HP-MA4 | 1:40 in 0.5%BSA/PBS |
| CD158b (KIR2DL2/DL3) | Miltenyi Biotec | 130-092-618 | Endometrial tissue | PE | DX27 | 1:20 in 0.5%BSA/PBS |
| CD3 | BD Biosciences | 644611 | Blood | FITC | SK7 | 20µL antibody mixture/50µL blood |
| CD16 | BD Biosciences | 644611 | Blood | PE | B73.1 | 20µL antibody mixture/50µL blood |
| CD45 | BD Biosciences | 644611 | Blood | PerCP-Cy5.5 | 2D1 | 20µL antibody mixture/50µL blood |
| CD56 | BD Biosciences | 644611 | Blood | PE | NCAM16.2 | 20µL antibody mixture/50µL blood |

